# Saccadic omission revisited: What saccade-induced smear looks like

**DOI:** 10.1101/2023.03.15.532538

**Authors:** Richard Schweitzer, Mara Doering, Thomas Seel, Jörg Raisch, Martin Rolfs

## Abstract

During active visual exploration, saccadic eye movements rapidly shift the visual image across the human retina. Although these high-speed shifts occur at a high rate and introduce considerable amounts of motion smear during natural vision, our perceptual experience is oblivious to it. This saccadic omission, however, does not entail that saccade-induced motion smear cannot be perceived in principle. Using tachistoscopic displays of natural scenes, we rendered saccade-induced smear highly conspicuous. By systematically manipulating peri-saccadic display durations we studied the dynamics of smear in a time-resolved manner, assessing identification performance of smeared scenes, as well as perceived smear amount and direction. Both measures showed distinctive, U-shaped time courses throughout the saccade, indicating that generation and reduction of perceived smear occurred during saccades. Moreover, low spatial frequencies and orientations parallel to the direction of the ongoing saccade were identified as the predominant visual features encoded in motion smear. We explain these findings using computational models that assume no more than saccadic velocity and human contrast sensitivity profiles, and present a motion-filter model capable of predicting observers’ perceived amount of smear based on their eyes’ trajectories, suggesting a direct link between perceptual and saccade dynamics. Replays of the visual consequences of saccades during fixation led to virtually identical results as actively making saccades, whereas the additional simulation of perisaccadic contrast suppression heavily reduced this similarity, providing strong evidence that no extra-retinal process was needed to explain our results. Saccadic omission of motion smear may be conceptualized as a parsimonious visual mechanism that emerges naturally from the interplay of retinal consequences of saccades and early visual processing.

## Introduction

Active observers sample their visual environment by making rapid eye movements – saccades. These frequent and staggeringly fast movements routinely induce drastic sensory consequences as they shift the entire image of the visual scene across the retina, yet we remain perceptually unaware of these changes. The phenomenon that we remain largely unaffected by the immediate consequences of our own saccades is one of the core problems of visual stability, known as saccadic omission. Despite decades of research in this area, first and foremost on the effect of saccadic suppression, that is, the reduction of contrast sensitivity around saccades (Binda & Morrone, 2018; E. Matin, 1974; Ross, Morrone, Goldberg, & Burr, 2001a; Volkmann, 1986), very little is known about the direct visual correlates of saccades, that is, intra-saccadic smear. Indeed, unless one is making eye movements across a small light source in a dark room – conditions under which elongated streaks are readily accessible (e.g., Bedell & Yang, 2001; E. Matin, Clymer, & Matin, 1972) – intra-saccadic smear is rarely observed in everyday life. Just as well, smear has hardly been investigated in the context of natural vision except for one prominent study: Campbell and Wurtz (1978) illuminated their laboratory by triggering a flash tube for 50-70 ms around the time of saccades, showing that flashes that occurred strictly during saccades resulted in a strikingly smeared appearance of the scene that “would be expected physically as the retina sweeps over the scene at a peak velocity of at least 500-600°/s” (p. 1298). Importantly, when the durations of flashes were extended beyond saccade duration, perceived smear rapidly dissipated and gave rise to a clear, unsmeared view of the scene. While many mechanisms contribute to saccadic omission at different stages of the visual hierarchy (for an overview, see, e.g., Rolfs & Schweitzer, 2022), the reduction of intensity due to smearing (Mitrani & Yakimoff, 1970, 1971) in combination with subsequent masking due to the presence of a post-saccadic retinal image seem to be sufficient to achieve perceptual omission of both intra-saccadic motion (Castet, Jeanjean, & Masson, 2002; Chekaluk & Llewellyn, 1990; Duyck, Wexler, Castet, & Collins, 2018) and motion streaks (Balsdon, Schweitzer, Watson, & Rolfs, 2018; Duyck, Collins, & Wexler, 2016; E. Matin et al., 1972; Schweitzer & Rolfs, 2020b).

Despite the relevance of this masking mechanism for the omission of intra-saccadic signals, it is surprisingly unstudied and many aspects of it remain elusive: What kind of visual information can remain resolvable during saccades? What is the subjective appearance of smear in natural environments, and what factors determine the consciously perceived extent of smearing? What is the relationship between intra-saccadic smearing and postsaccadic masking, and do these processes interact? And, finally, is the perception smear modulated by extra-retinal mechanisms, or can it be understood as a largely passive visual process? Understanding the nature of saccade-induced visual input could shed light on how the visual system achieves the impressive feat of omitting such input from conscious awareness, as well as whether processing that input could in principle contribute to visuomotor tasks or even visual stability.

In this paper we used a new tachistoscopic presentation technique that allowed the assessment of the dynamics of large-field intra-saccadic smear across a wide range of different natural scenes. Building on the basic finding of saccadic omission (Campbell & Wurtz, 1978), our goal was to understand the spatiotemporal transformations that the visual scene undergoes when it is rapidly shifted across the retina. We assessed the intra-saccadic time course of objective performance in smeared scene identification as well as the smear’s subjective appear-ance at a high temporal resolution. We furthermore investigated the impact of a variety of visual manipulations, such as spatial-frequency and orientation content, color, or viewing of windowed parts of a scene, and used computational modeling to synthesize the visual mechanisms that gave rise to the observed data. Ultimately, replaying the visual consequences of saccades during fixation, we experimentally validated the assumption that visual mechanisms alone could account for our findings. This effort provided new key insights about visual processing in the face of rapid eye movements. Specifically, our results suggest that saccade-induced smear can be characterized by a drastic redistribution of power towards orientations similar to the direction of the inducing saccade that varies contingently upon the saccadic velocity profile. Generation as well as omission of smear thus emerges as mere consequences of saccade dynamics and early visual processing, regardless of whether saccades are actively made or their visual effects are observed in a passive manner.

## Results

To study saccade-induced smear, we implemented an experimental paradigm that triggered tachistoscopic displays of natural scenes as soon as observers initiated saccades towards a remembered saccade target in near-complete darkness (Figure 1a). With this setup saccade-induced smear could be experienced as highly conspicuous, matching the initial observations made by Campbell and Wurtz (1978). Throughout three experiments we applied both an objective and a subjective task to assess observers’ perception of smear (see Task procedure in saccade experiments for details). In Experiments 1 and 2, observers performed a scene-matching task in which they simply indicated whether a scene presented during an eye movement was the same as a second scene presented after an interval of 500 ms when the eyes were fixating. In this setting, lower scene-matching performance indicated a larger impact of smear, as more strongly blurred scenes would be harder to identify. In Experiment 3, observers adjusted a dynamic motion filter to freely indicate their subjective experience of the previously presented scene. Here, the adjusted length of the motion filter was a direct measure of the amount of smear perceived during intrasaccadic presentation.

**Figure 1.**
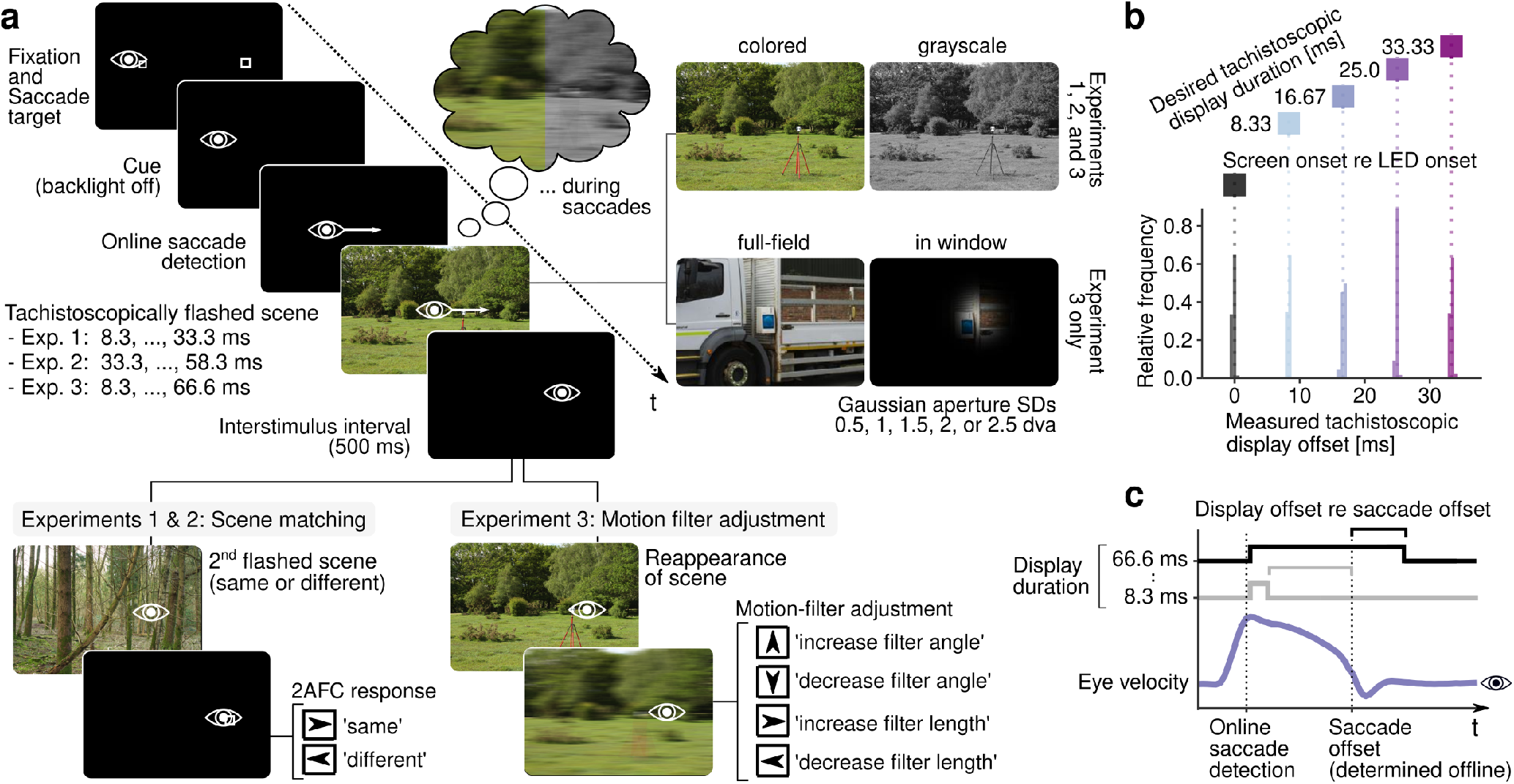
Experimental paradigm. **a** Task procedure used to investigate perception of intra-saccadic smear induced by natural scenes. Scene images, in either color or grayscale (all experiments), and Gaussian windows of the scene varied in size (Experiment 3) were flashed upon online saccade detection while the eye was still moving, resulting in a smear percept of the image. In Experiments 1 and 2 observers indicated whether the scene present during a saccade was the same as a second scene presented during fixation. In Experiment 3 observers adjusted a linear motion filter to match their subjective perception during the saccade. **b** Tachistoscopic display durations measured with a 2000-Hz lightmeter closely matched the specified display durations in Experiment 1. In addition, the latency of the display was as low, or even lower, than the latency of a single LED triggered by the same presentation schedule. **c** Time course of a presentation triggered by the saccade. To assess the notion of post-saccadic masking, we compute display offset relative to saccade offset. This metric quantifies the time that the illumination of the display was extended beyond the end of the saccade.

### Timing of tachistoscopic presentations

Using a tachistoscopic display in this paradigm had two major advantages compared to using a normal monitor. First, stimuli could be displayed gaze-contingently with much shorter latencies and increased temporal precision, because drawing stimuli to the screen while the monitor’s backlight was off saved an entire refresh cycle of the graphics card (Schweitzer & Rolfs, 2020a). Second, unlike CRT or LED monitors where pixels are known to have varying and possibly asymmetric rise and decay times (see, e.g., Komban et al., 2014), the tachistoscopic display activation profile is essentially a step function (see Tachistoscopic display for photodiode measurements). Results suggested not only that the pulse durations were accurate, but also that the display operated without considerable latency when compared to the latency of a single LED (Figure 1b). The presentation technique thus allowed a tight temporal synchronization of display on- and offsets with eye tracking data, for instance, when relating the time of display offset to the time of saccade offset (Figure 1c): While display onset relative to saccade onset was similar across experiments and duration conditions (Exp. 1: *F*(3, 57) = 1.91, *η*^2^ = 0.003, *p* = .138; Exp. 2: *F*(3, 57) = 1.25, *η*^2^ < 0.001, *p* = .299; Exp. 3: *F*(7, 133) = 0.39, *η*^2^ = 0.002, *p* = .904), display offset relative to saccade offset depended not only on display duration but also saccade duration – a variable impossible to control experimentally that was subject to considerable variability (Figure 2a, background histograms in middle and bottom panels). Display offset relative to saccade offset is the crucial variable as it captures the amount of time that visual input remains present beyond the end of the saccade and that therefore should determine the amount of post-saccadic masking (Campbell & Wurtz, 1978). To compute an accurate and sufficiently conservative estimate of saccade offset, which is (at least in the case of video-based eye tracking) prone to distortions due to postsaccadic oscillations (PSOs; Hooge, Holmqvist, & Nyström, 2016; Nyström & Holmqvist, 2010), we applied a re-cently proposed algorithm to detect onsets of PSOs (see Pre-processing for saccade experiments). PSOs were shown to coincide closely with saccade offsets measured by scleral search-coil techniques (Deubel & Bridgeman, 1995a; Schweitzer & Rolfs, 2022).

**Figure 2.**
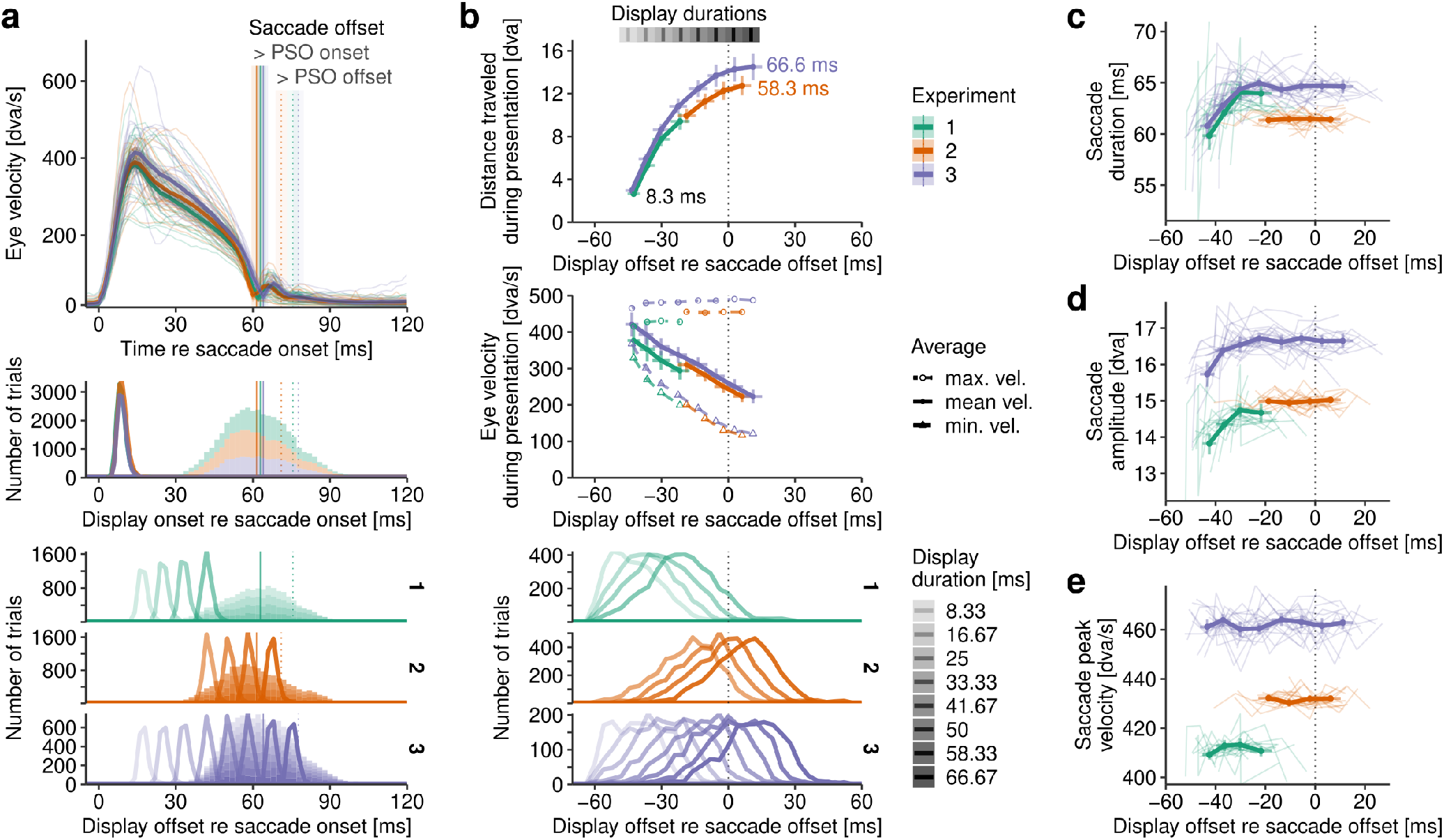
Timing around saccade offset. **a** Average saccadic velocity profiles (normalized to average duration; see also Figure A1) for each experiment (upper row). Vertical lines indicate saccade durations (solid: defined by the onset of the postsaccadic oscillation, PSO; dotted: defined by standard detection approaches). Below, frequency plots show distributions of display onsets (middle row) and offsets (bottom row) relative to saccade onset. Background histograms describe corresponding distributions of saccade durations in each experiment. **b** Average distance traveled (upper row) and velocity of the eye during the interval of display illumination (middle row) as a function of display duration and, as a consequence, display offset relative to saccade offset (bottom row). **c** Average saccade metrics are displayed for each available display duration in each experiment. Thin lines show averages of individual observers. Vertical and horizontal errorbars indicate ±1 standard error of the mean (SEM) of the dependent variable and ±2 SEM of the timing around saccade offset, respectively.

### Impact of tachistoscopic presentations on saccades

With respect to our experimental paradigm that did not manipulate display onset relative to saccade onset and the typical shape of the saccadic velocity (see average profiles in Figure 2a), three important implications should be noted (Figure 2b). First, the longer the display duration, the further the eye traveled while the screen was illuminated. If the amount of smear were a function of the size of the retinal shift, then it would be expected to be monotonically increasing. Second, the longer the display duration, the lower was the average velocity at which the eye traveled during the time of screen illumination. If smear increased with the overall velocity of the retinal shift, then one would expect its amount to monotonically decrease with longer display durations. Third, saccade durations introduced considerable variance in the timing of display offset around saccade offset. Importantly, we still grouped results by display duration to avoid confounding the dependent variables with any effect of varying saccade metrics. Note that computing aggregates from overlapping distributions with large variance (such as those shown in Figure 2b, bottom rows) could have heavily smoothed the time courses of dependent variables. To rule out this potential problem, we performed control analyses to directly investigate the impact of the saccade-duration variance on the temporal dynamics of perceptual judgments (Figure A2) but found no evidence that overall time courses were greatly altered when aggregates were computed on narrower saccade-duration distributions.

As all conditions and stimuli were presented in a randomly interleaved fashion, therefore making predictive adjustments of saccades unlikely, and visual latencies are typically considered too long to affect the motor command of the ongoing saccade, one would strongly assume that saccade metrics would be independent of display duration, that is, that the distributions of saccade metrics would be similar in each duration condition. Here we tested these assumptions using repeated-measures ANOVAs (see Statistical analyses of experimental results). Surprisingly, we found that this was not the case for short display durations: Saccade durations (Figure 2c) were shortened in both Experiments 1 (*F*(3, 57) = 12.67, *η*^2^ = 0.045, *p* < .001, *p*_*GG*_ < .001) and 3 (*F*(7, 133) = 12.76, *η*^2^ = 0.025, *p* < .001, *p*_*GG*_ < .001) but not in Experiment 2 (*F*(3, 57) = 0.06, *η*^2^ < 0.001, *p* = .980), and saccade amplitudes (Figure 2d) were reduced again in both Experiments 1 (*F*(3, 57) = 10.22, *η*^2^ = 0.036, *p* < .001, *p*_*GG*_ < .001) and 3 (*F*(7, 133) = 8.52, *η*^2^ = 0.013, *p* < .001, *p*_*GG*_ < .001) but not in Experiment 2 (*F*(3, 57) = 0.36, *η*^2^ < 0.001, *p* = .784). The finding that saccadic peak velocity (Figure 2e) remained unchanged in all experiments (Exp. 1: *F*(3, 57) = 2.22, *η*^2^ < 0.001, *p* = .095; Exp. 2: *F*(3, 57) = 1.10, *η*^2^ < 0.001, *p* = .354; Exp. 3: *F*(7, 133) = 1.05, *η*^2^ < 0.001, *p* = .399) suggests that saccades might have been stopped early when display-off transients occurred sufficiently early to still affect the ongoing movement. The inspection of individual saccadic velocity profiles (Figure A3) confirmed this suspicion, as mostly the decelerating section of saccades was affected, whereas the early, accelerating section remained similar across display durations. The profiles also put the effect of display duration on saccade duration in perspective: It was not directly observable in most observers and therefore of rather small size, that is, of 5 ms at most (Exp. 1: *M*_8.33_ = 59.84, *SD* = 6.27 vs. *M*_33.33_ = 63.96, *SD* = 9.77; Exp. 3: *M*_8.33_ = 60.78, *SD* = 6.52 vs. *M*_33.33_ = 64.97, *SD* = 8.98). It is therefore reasonable to assume that tachistoscopic presentations did not alter ongoing saccades to an extent large enough to systematically confound, or let alone explain, variations in perception of smear found throughout the course of the saccade.

### Time course of intra-saccadic smear perception

Experiments 1 and 2 surveyed to what extent scenes that were distorted by intra-saccadic motion blur could be matched to scenes that were briefly presented during fixation. Proportion of correct matches should therefore be inversely related to the amount of induced smear. We found that scene-matching performance followed a nonlinear time course (Figure 3a-c), similar to an inverted U, indicating that smear first increased in the first half of the saccade and then decreased in the second half. Note that this shape is highly consistent with previous studies having investigated the perceived length of intra-saccadic motion streaks (cf. Bedell & Yang, 2001; E. Matin et al., 1972).

**Figure 3.**
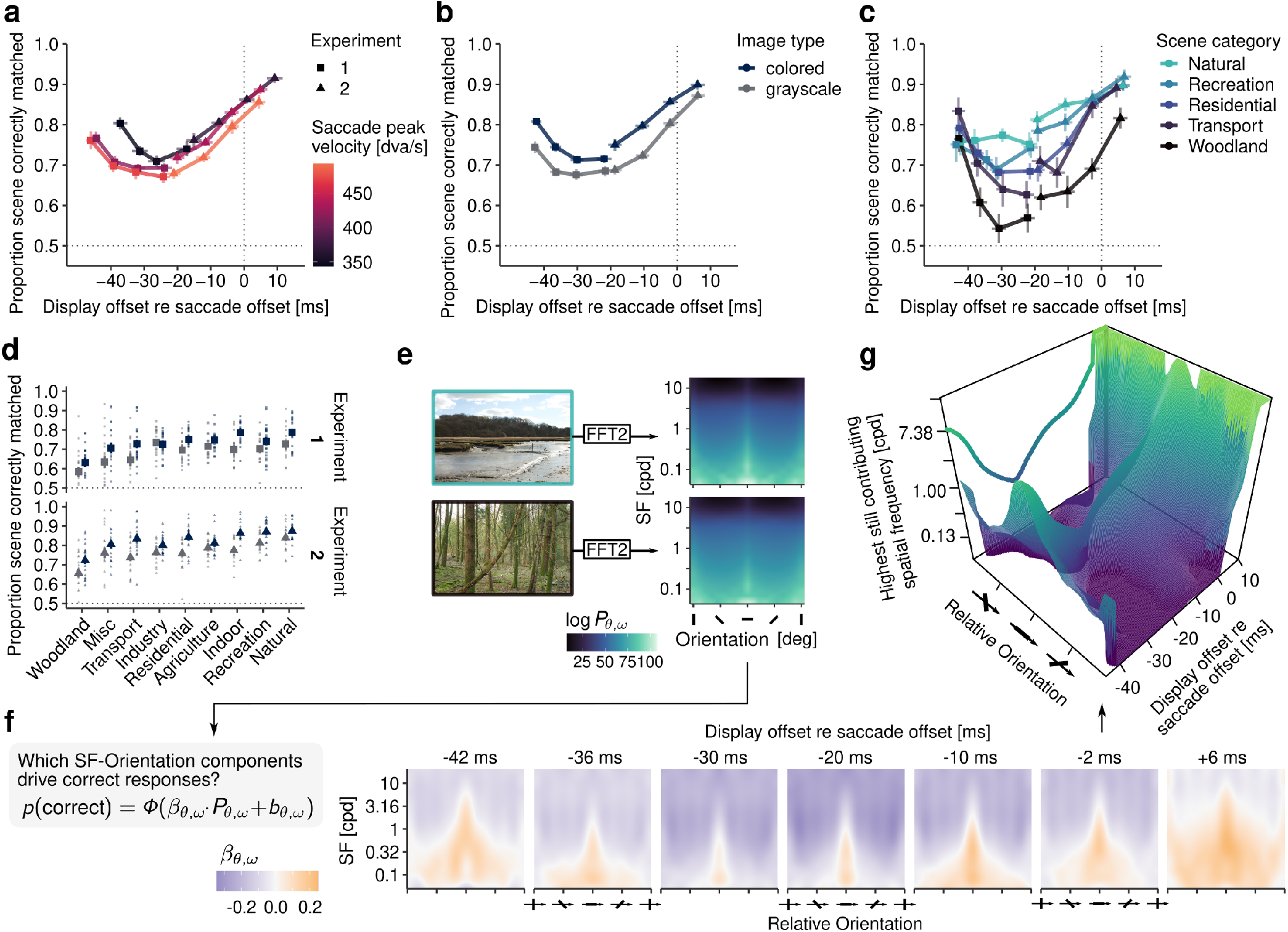
Scene-matching performance throughout the saccade (Experiments 1 and 2). **a-c** Proportion of correctly matched scenes as a function of display offset relative to saccade offset depending on saccadic peak velocity, image type, and scene category, respectively. Each point (rectangles: Experiment 1, triangles: Experiment 2) represents an experimentally manipulated display duration. Vertical error bars indicate ±1 within-subject SEM. **d** Sorted average task performance across display durations for colored and grayscale images in Experiments 1 and 2. Smaller dots indicate individual average performance. **e** Example stimuli from the natural-environment (upper) and woodland (lower) categories with their respective SF-orientation spectra. **f** Weights β_θ,ω_ indicate estimated slopes resulting from mixed-effects logistic reverseregressions fitted to predict correct responses from spectral power in individual SF-orientation components (see Reverse Regression). Each panel relates to one of the seven display durations. **g** 3D representation of the SF-orientation surface over time relative to saccade offset. The ordinate specifies the highest still contributing SF for a given relative orientation, i.e., the highest SF at which β_θ,ω_ ≥ 0.

In a first analysis, we investigated whether the instantaneous velocity of the ongoing saccade, around which display onsets occurred (see Figure 2a), had an impact on this time course (Figure 3a): We created three equal-sized bins based on the saccadic peak velocity distribution of each participant and found that, indeed, higher peak velocities resulted in lower task performance in both Experiment 1 (*F*(2, 38) = 6.02, *η*^2^ = 0.032, *p* = .005, *p*_*GG*_ = .009) and Experiment 2 (*F*(2, 38) = 25.52, *η*^2^ = 0.119, *p* < .001). Both experiments also showed the statistical signature of the time course, that is, significant main effects of display duration (Exp. 1: *F*(3, 57) = 14.37, *η*^2^ = 0.092, *p* < .001; Exp.2: *F*(3, 57) = 49.25, *η*^2^ = 0.379, *p* < .001, *p*_*GG*_ < .001) but no interaction between display duration and peak velocity (Exp. 1: *F*(6, 114) = 0.44, *η*^2^ = 0.004, *p* = .853; Exp.2: *F*(6, 114) = 0.40, *η*^2^ = 0.003, *p* = .874). Peak velocity thus had an impact on performance but not on the overall shape of the time course.

In a second analysis, we investigated the effect of color on scene-matching performance (Figure 3b). Color images were matched with clearly higher proportion correct in both experiments (Exp. 1: *F*(1, 19) = 47.02, *η*^2^ = 0.063, *p* < .001; Exp.2: *F*(1, 19) = 32.57, *η*^2^ = 0.111, *p* < .001). There was no interaction between image type and display duration in Experiment 1 (*F*(3, 57) = 1.14, *η*^2^ = 0.006, *p* = .337) and a small but significant interaction in Experiment 2 (*F*(3, 57) = 3.09, *η*^2^ = 0.013, *p* = .034, *p*_*GG*_ = .042), which suggests that the effect of color was largely additive but dissipated slightly when presentations were extended beyond saccade offset (Figure 3b). The effect of color may not be surprising, as color information is known to enhance the recognition of natural scenes displayed for short durations (Gegenfurtner & Rieger, 2000; Wichmann, Sharpe, & Gegenfurtner, 2002). In this context, it shows that the same benefit occurs for smeared scenes – possibly also because chromatic stimuli are less affected, although not unaffected, by saccadic suppression (Braun, Schütz, & Gegenfurtner, 2017; Knöll, Binda, Morrone, & Bremmer, 2011).

As human observers typically encounter a large variety of visual environments in their daily life, we also looked at the effect of scene category. In this third analysis, we computed observers’ task performance for each image category (as defined in the SYNS database by Adams et al., 2016) and display duration. Figure 3c shows the striking differences between five exemplary scene categories: Natural environments (e.g., beaches, heathland) were matched with high accuracy, whereas built environments and especially woodland scenes caused drastic reductions in task performance (Exp. 1: *F*(4, 76) = 10.93, *η*^2^ = 0.078, *p* < .001; Exp.2: *F*(4, 76) = 24.90, *η*^2^ = 0.182, *p* < .001). Differences between categories across display durations were similar in both experiments (Figure 3d) and were not driven by color information, as there was no significant interaction between category and color condition (Exp. 1: *F*(8, 152) = 0.90, *η*^2^ = 0.013, *p* = .517; Exp.2: *F*(8, 152) = 1.71, *η*^2^ = 0.017, *p* = .099). These results suggest that certain visual features, which may be prominent in some scenes categories but not others, must have been less affected by retinal motion and therefore enabled improved scene recognition.

### Description of smear in the domain of spatial frequency and orientation

Spatial frequency (SF) and orientation are the most basic features encoded in early vision. To determine which SF-orientation components drive high scene-matching performance, we extracted spectral power from all stimulus images (Figure 3e) and performed a reverse-regression analysis (see Reverse Regression) which used mixed-effects logistic regression to predict correct responses from log-power in each component. The sign and size of the fitted beta weight (*β*_*θ*,*ω*_) indicated whether high power in a given SF-orientation was beneficial or detrimental for task performance and to what extent. Figure 3f and Figure 3g show the results of this analysis fitted by a Generalized Additive Model (GAM; Wood, 2017), which was capable of closely approximating the data 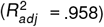.

Most importantly, high power around horizontal orientations – those parallel to the direction of the (mostly) horizontal saccades – were most beneficial for the correct identification of a scene (estimated degrees of freedom; *edf* = 10.91, *F* = 535.35, *p* < .001), especially at low SFs (*edf* = 97.51, *F* = 97.95, *p* < .001). In contrast, high power in oblique and vertical orientations – those orthogonal to saccade direction – were associated with more incorrect responses. This result is important but not necessarily surprising: Due to their high velocity, saccades induced high temporal frequencies (TFs) on the retina which would quickly render high-SF information unresolvable (Burr & Ross, 1982). For example, when viewing woodland scenes during saccades, the prominent vertical orientations in these scenes could not be perceived and caused observers to believe that a different scene was actually presented. In a similar vein, vertical gratings with a SF of 1.81 cpd were never perceived as moving when flashed during horizontal saccades of 6 degrees amplitude (Castet et al., 2002). In stark contrast, orientations parallel to their retinal trajectory remain resolvable during saccades, as they allow for more efficient temporal integration (Schweitzer & Rolfs, 2020b, 2021).

Moreover, the tuning around orientations parallel to the saccade’s direction, henceforth parallel orientations, followed a distinct time course across the saccade (Figure 3f), as confirmed by a significant three-way interaction term between orientation, SF, and display offset relative to saccade offset (*edf* = 169.96, *F* = 1.57, *p* < .001): At short display durations, which ended on average 42 ms prior to saccade offset, orthogonal orientations could still be used in the task provided SFs were small enough. The sharpest tuning occurred at intermediate display durations, reducing the set of useful SFs and orientation to a very small range around parallel orientations. Towards and after saccade offset, tuning again became less conspicuous as increasingly higher SFs and especially orientations orthogonal to saccade direction regained relevance. These orthogonal (i.e., vertical) orientations might have been especially relevant because they yield high power in man-made visual environments (Torralba & Oliva, 2003). These reverse-regression results already hint at a first tentative explanation for the time course found in our scene-matching task: At different time points throughout the saccade, observers may have had access to more or less restricted sets of the stimulus’ SF-orientation spectrum. A comparison of Figure 3f-g with Figure 3a-c provides evidence that a more limited band was indeed paralleled by a lower perceptual performance.

### Phenomenological appearance of smear

The analyses up to this point identified visual features that were most prominent in smeared scenes. The subjective appearance of smear, however, remained elusive. To shed light on the latter, Experiment 3 applied an adjustment task (see Task procedure in saccade experiments for details and supplementary video) that allowed observers to directly specify extent and direction of the experienced smear by setting the length and angle of a linear motion filter (Figure 1a).

Unlike the scene-matching task, this phenomenological approach produced responses that were directly related to the underlying perceptual judgment: The longer the adjusted motion filter length, the larger the amount of perceived smear (see Figure A6 for examples). The upper panel of Figure 4a shows adjusted motion filter length as a function of display duration, color, and full-field vs. window conditions. It becomes immediately evident that judgments strongly depended on display duration, which again determined the time of display offset (*F*(7, 133) = 17.33, *η*^2^ = 0.17, *p* < .001, *p*_*GG*_ < .001). Note that the time course is – if inverted – remarkably similar to the one found in Experiments 1 and 2 (cf. Figure 3b). Whereas this was the case for full-field scenes, time courses were significantly modulated when scene windows were shown (*F*(7, 133) = 21.81, *η*^2^ = 0.11, *p* < .001, *p*_*GG*_ < .001). Specifically, they were less steep compared to full-field scenes and quite similar to the masking time courses of (high-luminance) motion streaks measured by E. Matin et al. (1972), which also only peaked around 10-15 ms prior to saccade offset.

**Figure 4.**
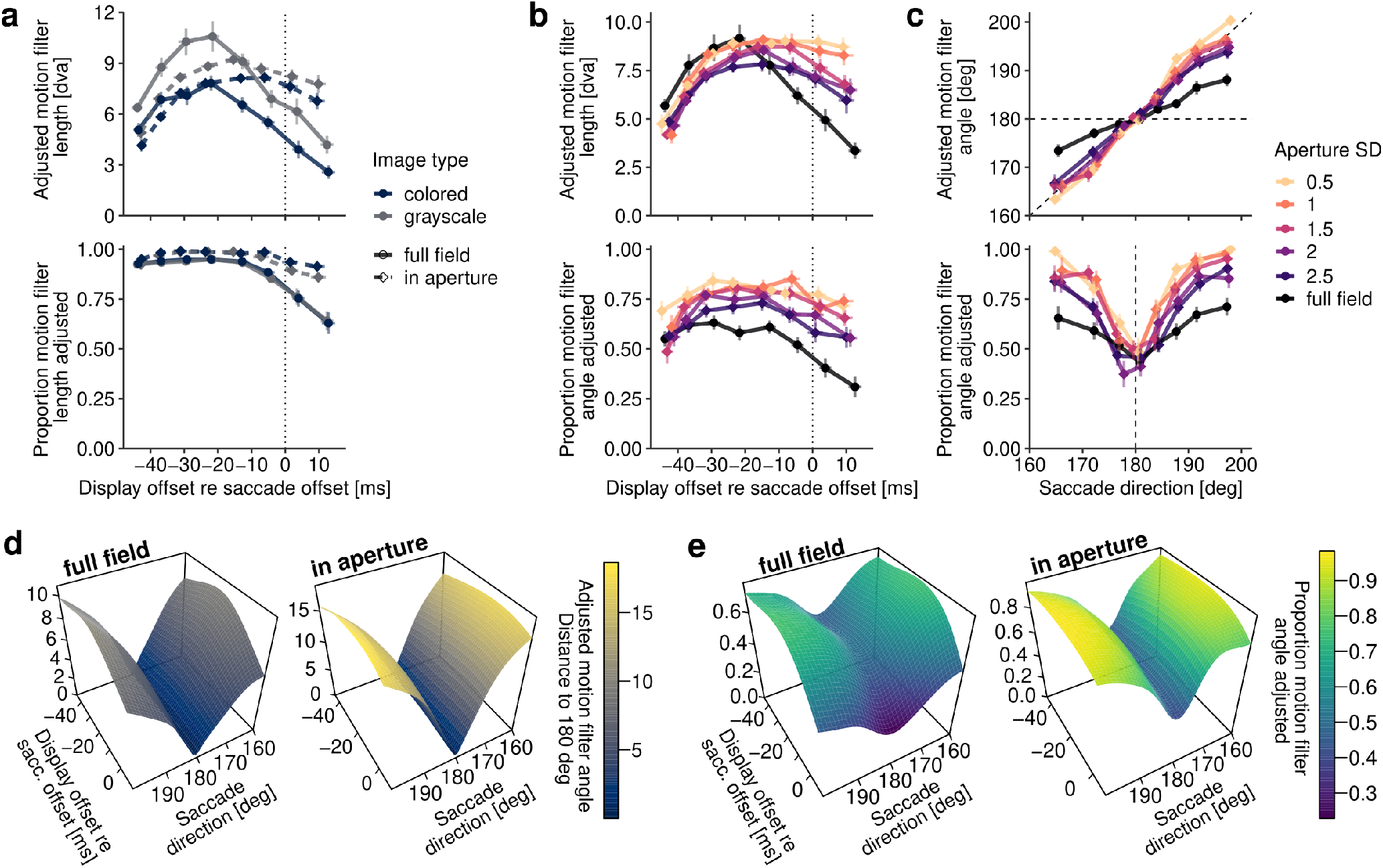
Subjective appearance of smear (Experiment 3). Responses were provided by adjusting the length and angle of a linear motion filter (for illustrations, see Figure A6). **a-b** Motion filter length adjusted (upper row) and proportion of adjustments (lower row, length and angle, respectively) as a function of display offset relative to saccade offset. **c** Motion filter angle adjusted (upper row) and proportion of angle adjustments as a function of saccade direction (8 bins per window SD condition). All vertical error bars indicate ±1 within-subject SEM. **d-e** Absolute distance of adjusted motion filter angles to horizontal (left) and proportion of angle adjustments (right) fitted by Generalized Additive Models incorporating predictors saccade direction and display offset relative to saccade offset.

In addition, a strong effect of color was found (*F*(1, 19) = 100.44, *η*^2^ = 0.06, *p* < .001), which was decreased when scene windows were shown (*F*(1, 19) = 17.55, *η*^2^ = 0.01, *p* < .001). Similar to Experiments 1 and 2, color did not affect the overall time course (*F*(7, 133) = 2.01, *η*^2^ = 0.004, *p* = .054, *p*_*GG*_ = .121). Observers adjusted motion filter length in a large proportion of trials (Figure 4a, lower panel). The proportion of adjustments however decreased towards and beyond saccade offset (*F*(7, 133) = 22.37, *η*^2^ = 0.21, *p* < .001, *p*_*GG*_ < .001), in particular for full-field presentations (*F*(7, 133) = 11.94, *η*^2^ = 0.07, *p* < .001, *p*_*GG*_ < .001). As display durations exceeded saccade durations, this indicates that observers more often perceived no smear at all but rather a sharp and static scene. This was less so the case for scenes presented in Gaussian windows, thus mirroring their much shallower post-saccadic masking time course. Note that this time course remained largely unchanged when individual condition means were computed based on narrower distributions of saccade duration (Figure A2c), suggesting that high variance in saccade durations did not strongly affect (i.e., smooth) time courses. As expected, sampling from a smaller range of time points slightly increased the steepness of time courses around saccade offset, but this very small effect was, if at all, restricted to full-field scenes.

We further looked at the adjustment of motion filter angles. As instructed saccade directions in Experiment 3 were varied within a range of 45 degrees, and saccade direction determined the direction of the shift induced on the retina, we specifically looked at the relationship of re-ported angle and saccade direction. Generally, observers adjusted motion filter angle less often than motion filter length (compare Figure 4a and Figure 4b, lower panels, respectively). As expected from previous results, angles were also adjusted more frequently when scenes were presented in windows (*F*(5, 95) = 18.09, *η*^2^ = 0.16, *p* < .001, *p*_*GG*_ < .001). When averaged across display durations, angle judgments were directly related to saccade direction (Figure 4c, upper panel). Slopes estimated by a linear mixed-effects model, in which factor window SD was treatment-coded, were close to 1 for the smallest window size (0.5 cpd: *β* = 1.17, *t* = 17.14, 95%CI [1.04, 1.31]) and decreased gradually, as window size increased (1 cpd: *β* = *−* 0.12, *t* = *−*2.41, 95%CI [-0.22, -0.02]; 1.5 cpd: *β* = *−* 0.20, *t* = *−*4.11, 95%CI [-0.29, -0.09]; 2 cpd: *β* = *−*0.23, *t* = *−* 4.77, 95%CI [-0.33, -0.14]; 2.5 cpd: *β* = *−*0.34, *t* = *−*6.99, 95%CI [-0.42, -0.23]). The largest decrease in slope was found for full-field scenes (*β* = *−* 0.72, *t* = *−* 14.96, 95%CI [-0.81, -0.64]), suggesting a strong bias to report smear as horizontally oriented, irrespective of the actual direction of the shift. This bias is likely driven by the strong prevalence of horizontal orientations in natural scenes (Coppola, Purves, McCoy, & Purves, 1998), which is also evident in the exemplary power spectra shown in Figure 3e. In window conditions, in contrast, the most prevalent orientation was the one introduced by the retinal trajectory of a single localized object, which appeared as a motion streak (Geisler, 1999; Geisler, Albrecht, Crane, & Stern, 2001; Jancke, 2000). This notion was further supported by the finding that the probability of adjusting the angle of motion filters depended on saccade direction – the more saccade directions deviated from horizontal, the more likely was the report of an angle (Figure 4c, lower panel). Theories of post-saccadic backward masking (Breitmeyer & Ganz, 1976; Castet, 2010; E. Matin, 1974) would to some extent predict that angle judgments follow a time course, as well: Orientations present in the stable post-saccadic scene should overrule orientations introduced due to retinal image motion. To investigate this question we fitted GAMs (see Statistical analyses of experimental results) to resolve these angle adjustments over time. The resulting surfaces (Figure 4d-e) indeed show that motion filter angles were reported less frequently and, if so, were increasingly perceived as horizontal when display offsets extended beyond saccade offset – even when saccade directions greatly deviated from horizontal. This suggests that saccadic omission not only reduced the extent of perceived smear but also diminished observers’ ability to report the direction of the image shift which the saccade undoubtedly must have induced.

### Spatiotemporal content of smear

Experimental results presented up to this point revealed that, first, perception of smear followed a distinctive continuous intra-saccadic time course which was largely similar in both scene-matching and adjustment tasks. Second, smear was heavily dominated by parallel orientations, that is, orientations parallel or close to the direction of the saccade. This was the case for the range of visual features that allow for efficient scene identification, as well as for motion filters that introduce power in those orientations specified by the filter’s direction.

Here we propose a model that attempts to explain these results in terms of early visual processes. For simplicity, this model (see Figure 5) considers a single (foveal) receptive field (RF) location that, as the eye is in midflight, is shifted across a static visual scene. During the shift, the RF received a sequence of input images over time, which were simulated based on the eye’s trajectory during the display of the stimulus image presented in any given trial in Experiment 3. We modeled RFs as Gabor filters with a given SF (0.05–5 cpd) and orientation (parallel to orthogonal orientations), thus manipulating the RFs SF-orientation-selectivity. Each RF, depending on its selectivity, was thus exposed to a certain luminance modulation (given by convolving the changing input image with the respective Gabor filter, see Spatiotemporal model) that it would respond to. It becomes immediately evident in Figure 5a that RFs selective to orthogonal orientations (i.e., on average vertical) and medium or even high SFs would respond strongly to the prevalent vertical orientations present in the scene. These orientations, however, also shifted through RFs at extremely high speeds and would therefore induce extremely high temporal frequencies (TFs). To confirm this suspicion, we computed TF power spectra (Figure 5b). Indeed, TF spectra exhibited prominent peaks whose position scaled with SF. For example, RFs orthogonal to the saccade’s direction, when exposed to 16.67 ms of visual stimulation (Figure 5b, top row, leftmost panel), received input of 525.8 Hz (*SE* = 21.6) at 1.58 cpd, 1014.2 Hz (*SE* = 48.7) at 2.81 cpd, and even 1781.4 Hz (*SE* = 79.5) at 5 cpd – a reasonable consequence given average eye velocities of close to 400 degrees per second (deg/s; Figure 2b). These peak TFs were reduced, but still extremely high, for oblique orientations (1.58 cpd: *M* = 338.3 Hz, *SE* = 17.8; 2.81 cpd: *M* = 709.8 Hz, *SE* = 35.6, 5 cpd: *M* = 1280.9 Hz, *SE* = 72.2), and practically absent for parallel orientations. Note that with increasing display durations and thereby later display offsets, peaks gradually lose their prominence, approaching a whitening across SFs in the domain of lower TFs (cf. Mostofi et al., 2020). Importantly, analyses up to this point did not consider the human contrast sensitivity profile (Figure 5c; see Contrast sensitiv-ity filters for details), and the fact that it is extremely unlikely that the visual system could resolve TFs as high the ones induced here during saccades. To determine the power of SFs and orientations in the resolvable range, we weighted spectral power according to temporal and spatial contrast sensitivity and summed across all available TFs. We found that sensitivity-weighted power in the middle-to-higher SF domain (i.e., SF ≥ 0.2 cpd) followed a specific time course that, when aggregated across SFs, looked remarkably similar to the time courses of task performance and extent of smear found in Experiments 1–2 and Experiment 3, respectively (Figure 5d). Low SFs barely showed any time course, as their peak sensitivity around 5–20 Hz was reached preferably during fast motion (Burr & Ross, 1982; Kelly, 1977; Robson, 1966).

**Figure 5.**
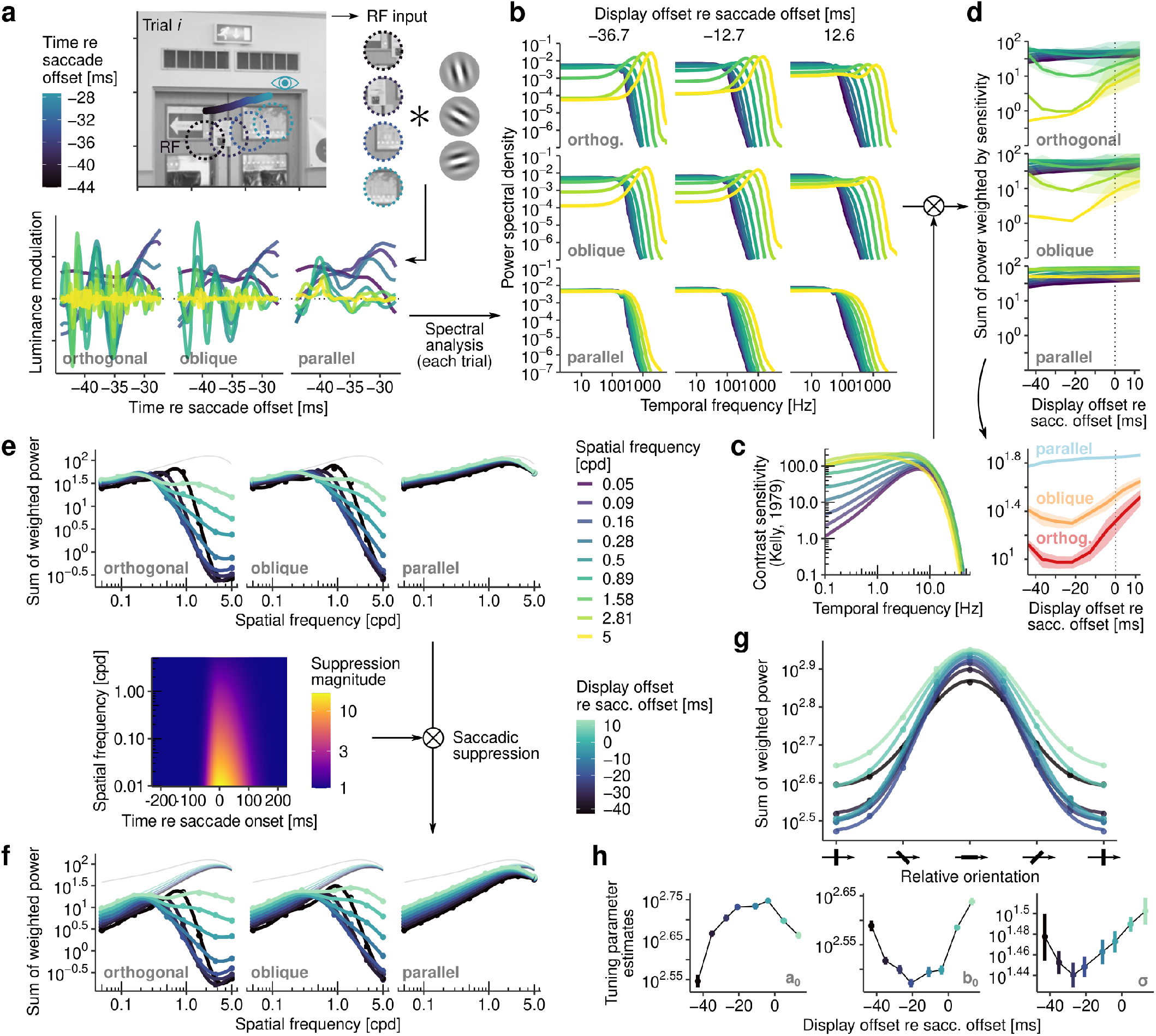
Spatiotemporal content of intra-saccadic smear. **a** Simulation of visual input to RFs, as they are shifted across the scene during saccades. Simulations were performed based on recorded saccade trajectories and stimulus presentations (see Spatiotemporal model), resulting in time-resolved Gabor filter responses for varying SFs (0.05–5 cpd) and orientations relative to the direction of the saccade, i.e., relative orientation. **b** Grand-average TF power spectra for all SFs, depending on relative orientation (rows) and display offset relative to saccade offset (columns). **c** Human contrast sensitivity profiles (based on Kelly, 1979) for available SFs and TFs indicating the range of resolvable information (see Contrast sensitivity filters). **d** Sum of power (across TFs) weighted by contrast sensitivity as a function of display offset relative to saccade offset. Bottom panel shows averages across SFs. **e-f** Sum of weighted power as a function of SF, before (top) and after (bottom) accounting for the effect of saccadic suppression (see Figure A4 for model details). Heatmap shows the magnitude of saccadic suppression depending on SF and time. **g-h** Sum of weighted power as a function of relative orientation, fitted by a Gaussian model. Fitted model parameters (bottom row) are shown as a function of display offset time.

Parallel orientations constitute a special case, as they remained largely unimpaired by the high velocity of the saccade. For example, when seen through an aperture, a grating moving in parallel to its orientation is not perceived as moving at all. Owing to the fact that a saccade never follows a perfectly straight path, a parallel grating is however never perfectly static and could therefore still be shifted in the TF range where contrast sensitivity is highest (Figure 5c). Note that this prevalence of parallel orientations mirrors previously described reverse-regression results, in which high power in these orientations predicted higher scene-matching performance (Figure 3f-g). Similarly, motion filters introduce high power in orientations parallel to the filters’ direction (for an illustration, see Figure A6). To further elaborate on this prevalence, we estimated orientation tuning functions using a more fine-grained orientation resolution (Figure 5g; for details see Spatiotemporal model). Unsurprisingly, all functions, regardless of the time of display offset relative to the saccade, peaked around orientations of zero (max(|*ω*_0_|) = 1.05^*°*^, *SE* = 0.28). All other tuning parameters (Figure 5h) systematically varied as a function of time. On the one hand, the amplitude parameter *a*_0_, indicating the difference between baseline and peak value, increased towards the end of the saccade before decreasing again. On the other hand, the baseline parameter *b*_0_ showed the inverse trend, indicating first a loss and then a recovery of power at orthogonal orientations. Finally, the tuning width *σ* corroborated these trends by exhibiting a time course again strikingly similar to behavioral findings: Similar to the reverse-regression results, tuning around parallel orientations was most narrow at display durations of 25 ms, that is, around 30 ms before saccade offset (cf. Figure 3f). Even though this model assumed nothing more than early visual SF-orientation selectivity, individual saccade dynamics, and human contrast sensitivity, it seemed to be well capable of capturing the prevalence of orientations parallel to saccade direction, as well as the conspic-uous time course found in experimental results.

### Effects of extra-retinal contrast suppression

The model can be described as purely visual, in other words, as not incorporating any potential extra-retinal modulations of visual sensitivity. Saccadic suppression describes the reduction of contrast sensitivity to stimuli presented around saccades that is not considered a mere consequence of visual changes induced by saccades (for a discussion see, e.g., Castet, Jeanjean, & Masson, 2001; Ross, Morrone, Goldberg, & Burr, 2001b). The effect is ubiquitous and also follows a well-documented time course contingent upon the execution of saccades (e.g., Diamond, Ross, & Morrone, 2000; Frost & Niemeier, 2015; Idrees, Baumann, Franke, Münch, & Hafed, 2020; Latour, 1962; Volkmann, Riggs, White, & Moore, 1978). One may thus ask if and, if so, to what extent the time course of smear perception could be explained by the dynamics at saccadic suppression. To this end, we modeled the time course (Figure A4a) and SF-specificity (Figure A4b) of saccadic suppression and adjusted contrast sensitivity profiles using the estimated magnitude of suppression at each corresponding time point and SF (Figure A4c-d). Figure 5e-f shows the resulting SF tuning for different display offset times before and after saccadic suppression. As described in the previous section, the high velocities of the saccade most strongly affected SFs ≥ 0.5 cpd, which is why these mid-high SFs were also subject to the greatest variation in power over time. In contrast, saccadic suppression is known to most strongly affect low SFs (Burr, Holt, Johnstone, & Ross, 1982; Burr, Morrone, & Ross, 1994; Niemeyer, Akers-Campbell, Gregoire, & Paradiso, 2022; Volkmann et al., 1978). Although this specificity to low SFs has been shown to depend to a considerable amount on the SF content of the visual environment (Idrees et al., 2020), it is quite reasonable to assume that it persisted in natural scenes which yield highest power in the low-SF domain (Tolhurst, Tadmor, & Chao, 1992; Van der Schaaf & van Hateren, 1996). Saccadic suppression thus decreased overall power, most strongly (i.e., up to ∼ 1 log unit) in the low-SF domain and shortly after saccade onset, but had little impact on the mid- or high-SF domain, which however exhibited the strongest modulation throughout the time course of the saccade. In other words, our model showed strongest effects at the higher spatial frequencies where the influence of saccadic suppression should be weakest – saccadic suppression was thus most likely not the cause of the observed results. Note also that time courses of saccadic suppression and time courses observed here differ in another respect: Saccadic suppression often peaks around the onset of saccades (for systematic reviews, see Binda & Morrone, 2018; Ross et al., 2001a; Volkmann, 1986), whereas both behavioral and modeling results here show lowest performance roughly around the midpoint of the saccade. While not being the cause of the observed effects, model results still seemed to suggest that saccadic suppression could have contributed to whitening across SFs by evening out the dominance of low-SF power (Figure 5f).

Recent evidence also suggested that saccadic suppression causes an enhancement of sensitivity in the high-SF domain that may start at as little as 2 cpd (Niemeyer et al., 2022). Although our model of SF-specificity only predicted an enhancement of SFs at approximately 7 cpd (Figure A4b), saccadic suppression may still have served to diminish the impairment of high SFs due to rapid image motion. It is however unlikely that this enhancement (i.e., a maximum enhancement of 80% at 5 cpd; Niemeyer et al., 2022) would be able to fully counteract an effect whose size is, at 5 cpd, of more than 2 log units in spectral power (Figure 5e).

### Saccade-induced bandpass filters

The model described in Figure 5 showed that large-field image motion due to saccades greatly constrains resolvable visual features in SF-orientation space. This bandpass filter is a direct consequence of human contrast sensitivity profiles and the direction and speed that a specific saccade imposes on the visual image at a given point in time. As illustrated in Figure 6a, when a scene is shifted through an aperture, the velocity of a certain orientation contained in the input depends on its angle relative to the direction of the shift. For example, an orientation orthogonal to the direction of movement would move at the same velocity as the saccade, whereas an orientation perfectly parallel to the direction of movement would not move at all. Saccades obviously induce high-velocity image shifts in the direction of their landing point, but they may also induce considerable velocities orthogonal to this direction due to saccade curvature (e.g., Smit & Van Gisbergen, 1990; Van der Stigchel, Meeter, & Theeuwes, 2006; Viviani, Berthoz, & Tracey, 1977). For instance, average peak velocity in the saccade’s direction amounted to 393.8 deg/s (SE = 16.1), whereas peak velocity orthogonal to the saccade’s direction amounted to 30.0 deg/s (SE = 4.3) (Figure 6b). This means that orientations parallel to the saccade’s direction are also considerably shifted within the RF’s aperture, and – depending on their SF – could therefore stimulate RFs in their ideal TF range (Figure 5c). Capitalizing on the fact that the velocity of individual orientations in a stimulus could be approximated at a given time throughout the saccade, we extended the contrast-sensitivity model by Kelly (1979) to construct SF-orientation bandpass filters (Figure 6c, see also Contrast sensitivity filters), whose effect is illustrated in Figure 6d. Right before the onset of saccades, when eye velocity was low, the contrast sensitivity profile peaked at mid-high SFs around 3 cpd (cf. Kelly, 1979, his Fig. 6) and was similar across all orientations, as both velocity components contribute equally. As saccades accelerated, peak sensitivity gradually shifted to lower SFs, as – while higher SFs became unresolvable – low SFs needed the saccade’s motion to reach the TF range where sensitivity would be highest. Parallel orientations were less affected by this shift, simply due to their reduced retinal velocity. Note that there was a high similarity between the shapes of resulting contrast sensitivity profiles (Figure 6c) and reverse-regression results (Figure 3f), not only in terms of the range of useful SFs and orientations around the peak velocity of the saccade but also in terms of how baseline sensitivity recovered towards and beyond saccade offset. As saccades decelerated, gradually higher SFs became resolvable again and the predominance of parallel orientations was alleviated, thereby reestablishing a clear and sharp image of the scene (Figure 6d).

**Figure 6.**
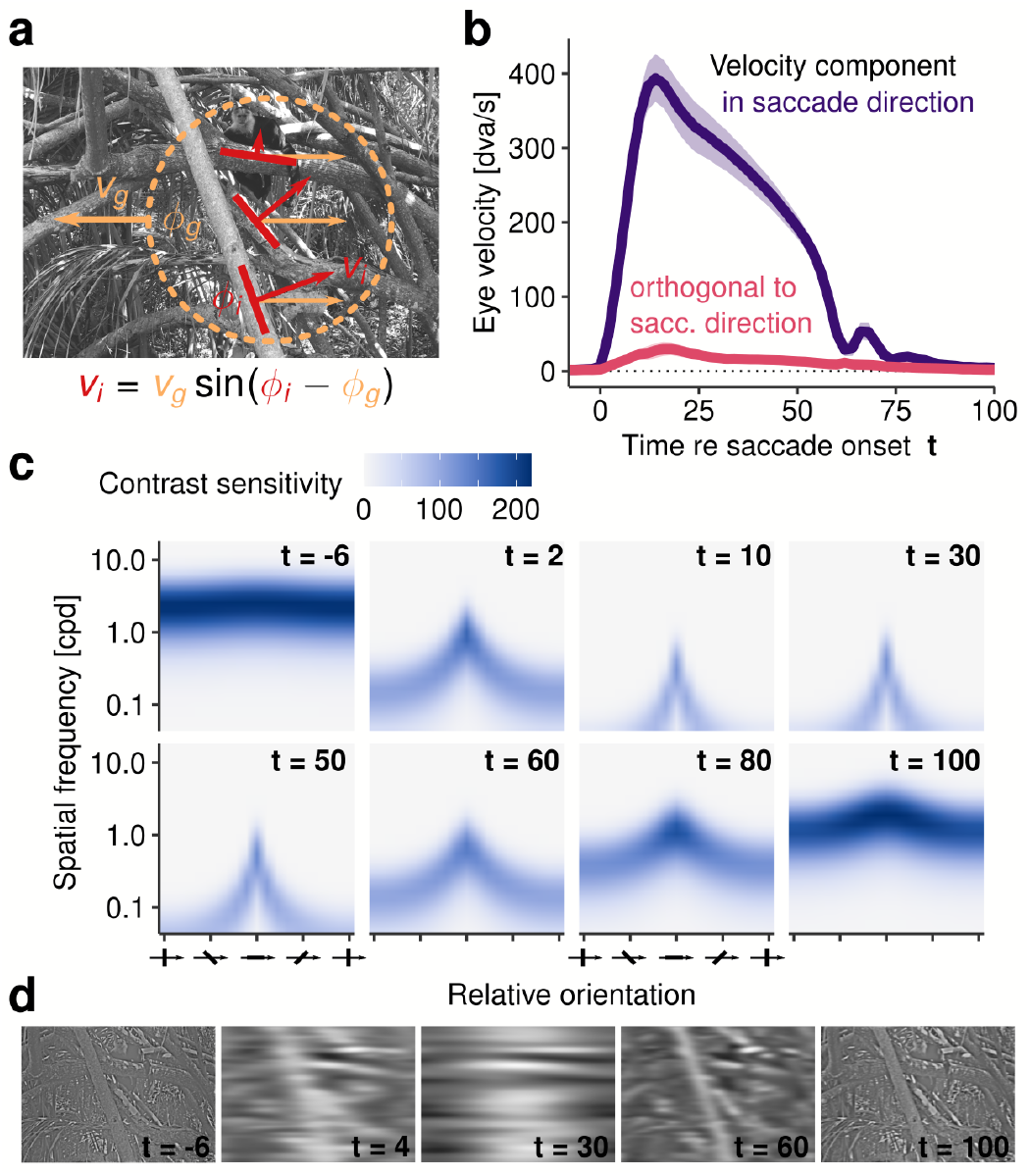
Smear as a velocity-dependent bandpass filter. **a** Illustration of the individual speeds v_i_ at which individual orientations ϕ_i_ travel as RFs are shifted across static visual scenes by saccades moving at a velocity v_g_ in a direction of ϕ_g_. **b** Mean absolute velocity of saccades over time relative to the onset of each saccade. Velocity was separated in two components, that is, in and orthogonal to the saccade’s direction, respectively. The small peak around 65-70 ms indicates the post-saccadic oscillation (cf. Figure 2a). **c** Contrast sensitivity profiles at different time points throughout the saccade (see Contrast sensitivity filters for model details) resulting from the average saccade profile shown in panel b. **d** Scene image (from panel a) bandpass-filtered according to the contrast sensitivity profiles at five selected time points, as shown in panel c.

### The motion-filter model of saccadic smear

To think of saccade-induced smear in terms of velocity-dependent SF-orientation bandpass filters may well describe what visual features can remain perceivable during saccades made across natural scenes but leaves out one important aspect that should greatly determine the amount of perceived smear, that is, the extent of the retinal shift. Campbell and Wurtz (1978) noted that when presentations were “left on for a duration of 5 msec or less […] the room was seen clearly no matter where in the 50–70 msec eye movement period the period of light occurred” (p. 1298). Indeed, even if the eye moved at high velocities, at extremely short presentation durations the extent of the retinal shift would be short enough to induce no visible smear whatsoever. This intuition is confirmed by the finding that the shortest presentation durations used here (i.e., 8.33 ms), even though these presentations were long enough to induce visible smear and occurred during the highest saccadic velocities (Figure 2b), elicited neither the lowest perceptual performance in Experiment 1–2 (Figure 3a-c) nor the maximum amount of perceived smear in Experiment 3 (Figure 4a-b) – most likely because the image shifted rarely more than 3 degrees of visual angle (dva; Exp. 1: *M* = 2.64 dva, *SE* = 0.13; Exp. 3: *M* = 2.94 dva, *SE* = 0.11) during that time. Therefore, the simplest hypothesis – without even assuming any properties of the visual system, such as a contrast sensitivity profile – would be that the amount of (perceived) smear is a function of the distance that the eye travels across a static scene.

We illustrate this idea in Figure 7a: We considered a RF that received varying inputs over time and integrated them, assigning higher weights to more recent input (for details, see Figure A5). This simple procedure inevitably not only produced smear but also masked it when largely constant post-saccadic inputs overweighted highly variable intrasaccadic inputs. Importantly, this model, which averaged pixel values that lie along a defined trajectory, could be generalized to a motion filter: Motion-filtered images, such as the ones adjusted in Experiment 3 (see examples in Figure A6), were created by convolving the scene image with a kernel matrix describing a linear trajectory (Figure 7b, left panel). Although motion-filtered scenes were sometimes experienced as strikingly similar to perceived smear, a linear trajectory is hardly a suitable model for smear induced by saccadic eye movements for three reasons. First, saccades do not move at a constant velocity but follow an approximately gamma-shaped velocity profile (Van Opstal & Van Gisbergen, 1987). Second, saccades do not follow a straight line but can exhibit considerable curvature (Figure 6b). Third, temporal dynamics of visual processing (e.g., Figure A5a) make it unlikely that all input, regardless of its recency, is weighted equally. To overcome these limitations, we created trajectory-dependent motion-filter kernel matrices (Figure 7b, right panel) that produced motion-filtered images according to each trial’s saccadic profile (for method details, see Motion filters).

**Figure 7.**
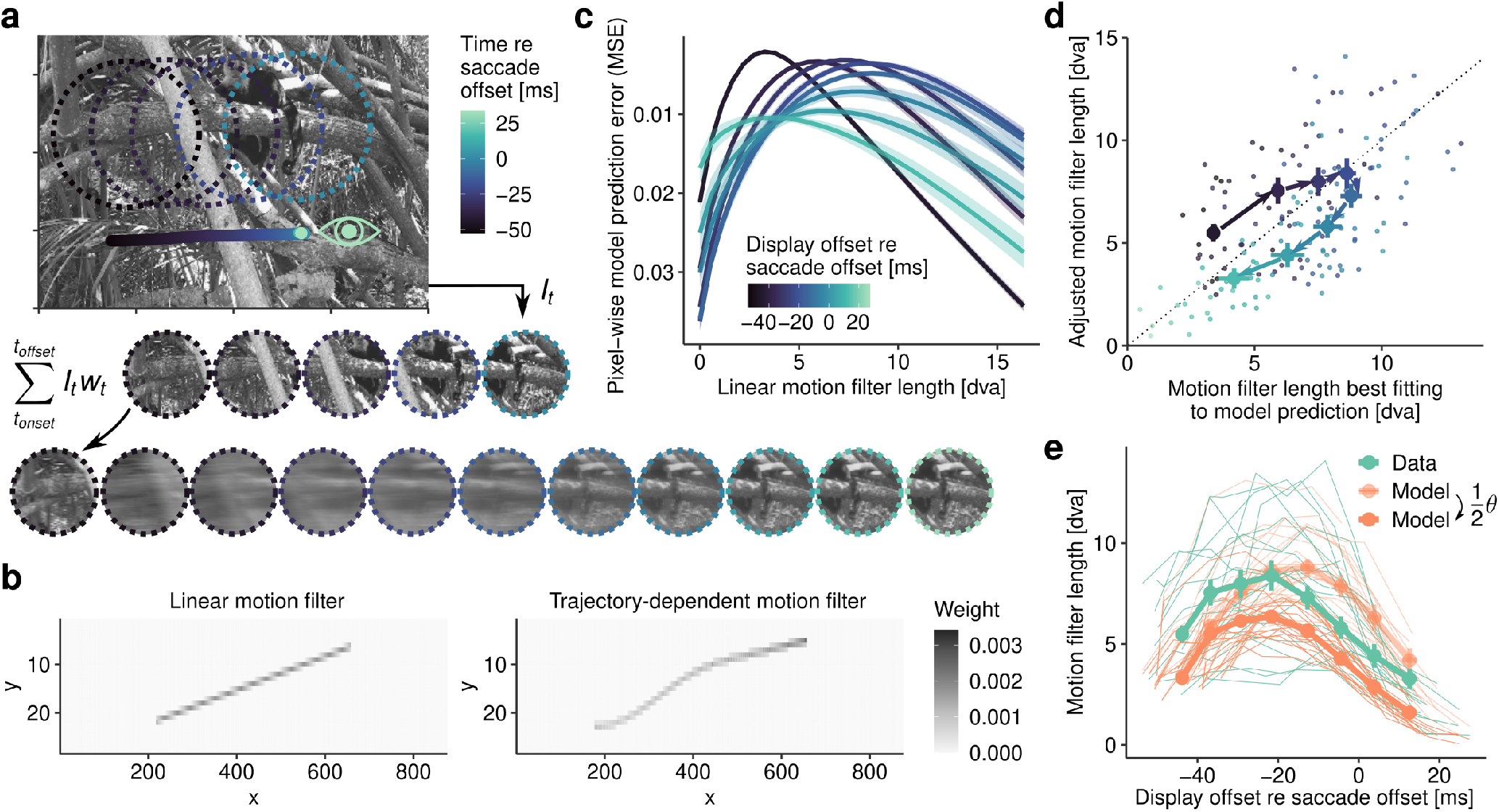
Smear as a motion-blur filter. **a** Illustration of a RF that is shifted across a scene by a saccade. Inputs I_t_ are integrated over time by computing a weighted sum, where the weight w_t_ depends on the recency of the input (Motion filters). Bottom row shows results simulating display durations from 1 to 100 ms (left to right). **b** Example motion-filter kernel matrices for a standard linear (left) and a custom trajectory-dependent motion filter (right), which was created based on measured gazeposition samples and a model of visual decay (Figure A5). **c** Curves indicate pixel-wise similarity (measured by mean square error, MSE) between images resulting from trajectory-dependent motion filters and comparison images resulting from linear motion filters of varying lengths. Each curve corresponds to one display duration. Shaded areas correspond to ± 1SEM. **d** Relationship between predictions of trajectory-dependent motion-filter model and linear motion filter length adjusted by observers. Small dots indicate individual observer means for each display duration, whereas larger dots indicate means across observers. All error bars indicate ± 1 SEM. Color coding is the same as in panel c. **e** The effect of the temporal response function on the time course estimated by the motion-filter model. Transparent orange line describes model predictions assuming an empirically estimated scale parameter of θ = 20.94 (see Figure A5a), whereas thick orange line describes model predictions assuming half this scale, that is, θ = 10.47. Data, shown in blue, represents the adjusted motion filter lengths (Experiment 3) for full-field scenes, averaged across color conditions (cf. Figure 4a).

To evaluate how well such a dynamic motion filter could approximate the linear motion-filter adjustments that observers provided in a given trial, we first computed how similar (quantified by mean square error, MSE) the image created by our trajectory-dependent motion-filter model was compared to a range of images created by linear motion filters of varying lengths. Figure 7c shows the mean similarity curves that resulted from this analysis. Similarity curves greatly varied over display offsets relative to saccade offset, with their peaks shifting to longer filter lengths before gradually shifting back towards zero, thereby approaching the original, unfiltered image. Overall similarity decreased over time, which is unsurprising, as trajectory-dependent and linear filter kernels became increasingly different. For example, a small portion of the saccade may still be approximately linear, but a filter incorporating the entire saccade trajectory will likely be far from linear. As a second step, we extracted each trial’s linear motion filter length at which similarity was largest, so that they could be compared to the motion filter length that observers reported in Experiment 3.

This comparison, revealed a moderately strong correlation between predictions made by the trajectory-dependent motion-filter model and observers’ given response (Figure 7d; *r* (158) = .60, 95% CI [0.49, 0.69], *p* < .001). Testing against a linear relationship with a slope of one and an intercept of zero, there was considerable variance around the unity relationship (*R*^2^ = 0.28), suggesting that the model only loosely predicted the absolute value of observers’ responses. Across display durations, the model was reasonably accurate (*M* = − 0.29 dva, *SE* = 0.59) but showed considerable absolute errors (*M* = 2.12 dva, *SE* = 0.36), which were quite systematic. First, the model underestimated the amount of perceived smear in the first half of the saccade by up to 2.1 dva (*SE* = 0.44). As the model was constrained by the extent of the retinal shift, its predictions were necessarily very close to the actual distance traveled by the eye during presentation (8.33: *M*_*dist*_ = 2.94 dva, *SE* = 0.11, *M*_*pred*_ = 3.38 dva, *SE* = 0.13; 16.67: *M*_*dist*_ = 5.91 dva, *SE* = 0.21, *M*_*pred*_ = 5.92 dva, *SE* = 0.24; cf. Figure 2b). It may thus be more appropriate to assume that observers overestimated the amount of smear produced by the retinal shift, which was, interestingly, less pronounced when they viewed windows of scenes (Figure 4a-b). Second, the model overestimated the amount of perceived smear shortly before and after saccade offset by up to 2.06 dva (*SE* = 0.38), in other words, it underestimated how rapidly the perceived smear was reduced around saccade offset. In our model this time course of smear reduction is governed by the sluggishness of the assumed temporal re-sponse function (Figure A5) – the more sluggish the response, the stronger the persistence of responses to intrasaccadic input in the system. To show this effect, we reduced the scale parameter of the temporal response function to 50% of its initial value. Indeed, the model now produced a time course strikingly similar to the empirically observed time course (Figure 7e), greatly increasing correlations with observer’s responses (*r* (158) = 0.70, 95% CI [0.61, 0.77], *p* < .001). To validate these results, we fitted a linear mixed-effects model that revealed estimated slopes close to 1 (*β* = 0.94, *t* = 7.61, 95% CI [0.68, 1.18]) but with significantly non-zero intercepts (*β* = 2.19, *t* = 5.40, 95% CI [1.39, 3.03]). This indicates that the model almost perfectly reproduced the time course of observed data, while being (by definition) unable to match observers’ higher baseline in the estimation of smear. In addition, it may have been possible that computing pixel-wise image similarity did not capture the perceptual decision-making of observers in Experiment 3 – indeed, it is known that *𝓁*_2_-based metrics are inherently dissimilar to human similarity judgements, especially in the presence of blur (Zhang, Isola, Efros, Shechtman, & Wang, 2018). To provide a more intuitive display of the model’s predictions, we plotted images filtered by the trajectory-dependent motion filter across all eight display durations in Figure A7. To conclude, a considerable extent of the overall shape of the time course observed in Experiment 3 could be predicted based on the saccade’s trajectory alone, thereby establishing a direct link between the saccade-induced retinal shift and the perceived amount of smear.

### Smear induced by simulated saccades

The models up to this point all have one thing in common: To explain generation and reduction of smear, they assume no more than the spatiotemporal properties of the stimulus (i.e., SF-content and time-resolved trajectory of the stimulus) and the spatiotemporal properties of the observer’s visual system (i.e., contrast sensitivity and visual decay). If these assumptions are valid, we can make a strong prediction: As long as the spatiotemporal dynamics in stimulus and observer remain, the presentation of a saccade’s visual consequences to the fixating eye should lead to the same experience of smear as when a saccade is actively executed across a static image.

Experiment 4 tested this critical hypothesis. Observers performed a task identical to the one used in Experiments 1 and 2 (Figure 8a), with one crucial difference: Instead of making a saccade to trigger the first scene presentation, observers viewed – during fixation – a replay of a trial sampled from saccade experiments. This replay comprised the exact same visual scene, display timing, and saccade trajectory (see Simulated saccades for details), replayed in retinotopic coordinates using a projection system with sub-millisecond precision. While such a stimulus would match the retinal dynamics of saccade-induced motion, we went a step further to additionally simulate the extraretinal dynamics of saccadic suppression, that is, the reduction of contrast sensitivity around saccades. Using the predictions of the descriptive Saccadic suppression model proposed in this paper, we filtered scene stimuli according to the suppression profile at each time point during presentation (see example in Figure 8b and, for details, Simulated saccadic suppression), thereby replicating not only the spatial but also the temporal pattern of saccadic suppression. If saccadic suppression, as a mechanism to dampen visual sensitivity to low SFs and motion during saccades (Binda & Morrone, 2022; Ross et al., 2001a), could explain (or at least contributed to) the results of Experiments 1 and 2, then one would expect that replays with combined retinal motion and suppression (henceforth motion+suppression replays) would more closely approximate the patterns of results found in saccade experiments than retinal motion alone.

**Figure 8.**
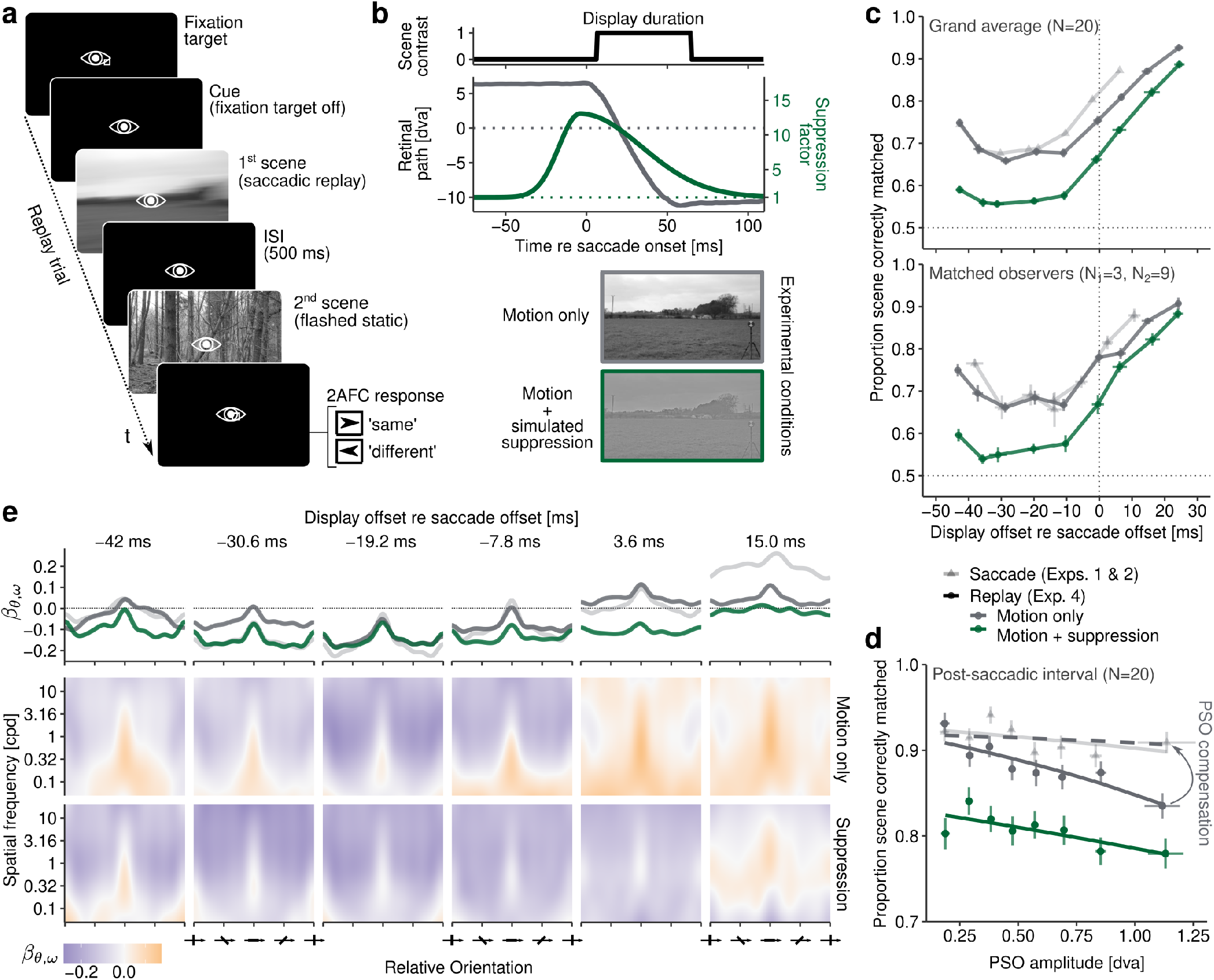
Perception of smear during simulated saccades (Experiment 4). **a** Task procedure to investigate smear induced by replayed saccades during fixation at 1440 fps. During replay trials, observers were instructed to fixate while performing a 2AFC, same-different scene-matching task, identical to Experiments 1 and 2. The first scene was presented moving according to the exact trajectories and stimulus timings recorded in those experiments (see Simulated saccades), whereas the second scene was static. **b** In addition to simulation of retinal motion, we used the Saccadic suppression model described in this paper to simulate the impairment of contrast sensitivity induced by saccadic suppression, that is, by filtering the image in a time-resolved manner (see Simulated saccadic suppression). The curve displays an exemplary suppression time course for SFs around 0.02 cpd. The scenes below illustrate the visual effects of filtering at the time of peak suppression. **c** Proportion correct in the scene-matching task as a function of display offset relative to saccade offset across all observers (top) and for observers who also participated in Experiments 1 and 2 (bottom). Results from saccade experiments are shown as transparent gray lines. Error bars indicate ±1 SEM. **d** Reverse-regression results with the same conventions as results from saccade experiments (Figure 3) with the top row showing the marginal average β weights across SFs. **e** Task performance (for trials with post-saccadic offset only) as a linear function of the amplitude of the post-saccadic oscillation (PSO) in the replay conditions and the saccade experiments (transparent gray line), respectively. The dashed line indicates the theoretical performance in the critical replay condition if PSOs had been displayed that would match the actual perceptual consequences of ocular lens shifts (see Estimation of perceptual consequences of post-saccadic oscillations).

Figure 8c shows the results of Experiment 4. First, it becomes immediately evident that simulated saccades, too, were capable of producing the prominent U-shaped time course we previously found in saccade experiments (*F*(8, 152) = 215.65, *η*^2^ = 0.63, *p* < .001, *p*_*GG*_ < .001). Second, it is equally striking that, compared to the simulation of saccadic motion alone, combined motion and suppression led to a drastic decrease in accuracy (*F*(1, 19) = 354.73, *η*^2^ = 0.27, *p* < .001), leading to intra-saccadic task performance near chance level during scene movement (*M* = 0.57, 95% CI [0.54, 0.59]). These results were underlined by the subjective appearance of these trials: Observers very consistently reported a gray-out, a near-complete loss of contours or discernible information, making it nearly impossible to solve the task. In accordance with the reports of Campbell and Wurtz (1978), however, trials without simulated suppression led to large amounts of smear, pushing task performance well above chance level (*M* = 0.69, 95% CI [0.65, 0.73]). As predicted by the time course of saccadic suppression (Figure 8b), the suppression manipulation yielded the strongest effects in the early phases of the saccade, whereas towards and beyond saccade offset the difference between the two experimental conditions was gradually reduced (*F*(8, 152) = 7.00, *η*^2^ = 0.05, *p* < .001, *p*_*GG*_ < .001).

Even though time courses in saccade and replay experiments were remarkably similar (Figure 8c, upper panel), especially in the early intra-saccadic interval, a direct statistical comparison between experiments is less straightforward. We therefore retrospectively matched observers across experiments based on their pseudonymized participant codes: Three observers participated in both Experiments 1 and 4, and nine observers participated in both Experiments 2 and 4. For these subsets, displayed in Figure 8c (lower panel), we could thus perform within-subject hypothesis testing. On the one hand, testing saccade vs motion-only replay as a within-subject condition – even though the respective experiments were conducted with a time lag of more than two years – revealed no average performance differences, neither in (intra-saccadic) Experiment 1 (*F*(1, 2) = 0.10, *η*^2^ < 0.01, *p* = .782) nor in (post-saccadic) Experiment 2 (*F*(1, 8) = 1.17, *η*^2^ < 0.01, *p* = .311). On the other hand, comparing saccades with the motion+suppression replay resulted in clear performance differences to both Experiment 1 (*F*(1, 2) = 1.89, *η*^2^ = 0.80, *p* = .005) and 2 (*F*(1, 8) = 52.70, *η*^2^ = 0.29, *p* < .001). Apart from these average performance differences, we also found no interactions between viewing condition and display durations (Exp. 1, motion-only: *F*(3, 6) = 0.66, *η*^2^ = 0.12, *p* = .602, *p*_*GG*_ = .507; Exp. 1, motion+suppression: *F*(3, 6) = 0.10, *η*^2^ = 0.22, *p* = .461, *p*_*GG*_ = .440; Exp. 2, motion+suppression: *F*(3, 24) = 0.64, *η*^2^ = 0.01, *p* = .591, *p*_*GG*_ = .553), suggesting that the shape of time courses for saccade and replay experiments were also largely similar, both in motion-only and motion+suppression replays.

Yet, there was one interesting exception that can be spotted in Figure 8c (upper panel): When comparing saccades with motion-only replay, the early postsaccadic interval exhibited significant interaction, indicating lower performance during replay than during saccades (*M*_*sacc*,58.33_ = 0.89, *M*_*replay*,58.33_ = 0.83; *F*(3, 24) = 4.74, *η*^2^ = 0.04, *p* = .010, *p*_*GG*_ = .012). Results

up to this point suggest a high similarity between saccade and motion-only replay experiments, whereas motion+suppression replays produce significantly lower task performance than saccades. The small but significant post-saccadic difference between saccades and motion-only replays merits further attention. Whereas such difference could be suspected to be related to some form of post-saccadic enhancement (Ibbotson, Price, Crowder, Ono, & Mustari, 2007), we hypothesized that it originated from post-saccadic oscillations (PSOs), which we left intact in replayed saccades. Specifically, PSOs arise as artifacts of video-based eye tracking, that is, the relative movement of pupil and corneal-reflection signals (for a review, see Schweitzer & Rolfs, 2022). Such PSOs routinely reach amplitudes of 0.5 dva and speeds of 50 dva/s, and can thereby introduce considerable motion signals when used for replay of saccade trajectories. Indeed, as shown in Figure 8d, PSOs had a considerable impact on postsaccadic task performance in the replay experiment: Using (treatment-coded) multiple logistic regression fitted on single-trial data, we estimated a reduction of the odds of a correct response by 0.49 per degree of PSO amplitude in the motion-only replay (*β* = −0.71, *t* = − 4.77, 95% CI [-0.99, -0.41], *p* < .001). Relative to this, the slope for motion+suppression replays was less steep (*β* = 0.40, *t* = 2.07, 95% CI [0.021, 0.78], *p* = .038), which was expected given the reduced power in relevant SF bands. Unlike in replay conditions, post-saccadic task performance in saccade experiments exhibited, if at all, a marginally significant dependence on PSO amplitude (*β* = −0.31, *t* = −1.92, 95% CI [-0.63, 0.013], *p* = .055). Yet, estimated regression intercepts for replay (*β* = 2.43, 95% CI [2.22, 2.63]) and saccade experiments (*β* = 2.54, 95% CI [2.32, 2.75]) were highly similar. This last result seems to suggest that in the absence of PSOs both saccade and replay conditions should produce largely the same task performance. Since PSO-induced retinal shifts arise from an angular offset of the lens (Deubel & Bridgeman, 1995a; Tabernero & Artal, 2014) and their perceptual consequence is much smaller than the measurable PSO amplitude itself (Deubel & Bridgeman, 1995b), it is likely that the replay experiment greatly overestimated the amount of post-saccadic motion effectively perceived, thus producing reduced task performance around saccade offset. To confirm this notion, we compensated for the effect of PSOs on task performance in the replay condition by converting the PSO amplitude we measured with P-CR eye tracking to the size of the perceptual shift that should arise from it (for details, see Estimation of perceptual consequences of post-saccadic oscillations). After applying this conversion to our regression results, task performance in the motion-only replay condition tightly approximated task performance in the saccade condition (Figure 8d), providing strong evidence that the post-saccadic difference between saccade and replay could be almost fully attributed to PSOs.

Finally, one may wonder whether the description of smear in the domain of SF and orientation has an equally high similarity between saccade and replay as their respective time courses, and how simulated saccadic suppression affects what SF-orientation bands can be used to perform scene matching. To that end, we performed the same reverse-regression analysis on replay results as previously on saccade results (for methodological details, see Reverse Regression). Reverse regressions revealed a strikingly high visual similarity between the SF-orientation patterns of replays (Figure 8e) and saccades (Figure 3f). Like saccade patterns, both replay patterns (with and without suppression) exhibited clear tuning around orientations parallel to saccade direction (see marginal weights in top panel of Figure 8e), as well as a time course with the narrowest SF-orientation band at around 30-20 ms before saccade offset. As predicted by our velocity-dependent bandpass-filter model (Figure 6), this shows that the dynamics of retinal motion alone could explain the emergence of these defining features of smear even when no saccade was executed. At closer inspection, it becomes evident that patterns resulting from motion-only replays, compared to motion+suppression replays, are more similar to those resulting from saccades. To quantify this impression, we computed correlations between individual patterns and found that motion-only patterns were better correlated to saccade patterns (*d* = 0.072, 95% CI [0.014, 0.130], *t* (5) = 3.20, *p* = .024). To better approximate the similarity between patterns, without taking into account absolute weights differences, we thresholded the data, selecting only those SF-orientation components that drove correct responses (i.e., *β*_*θ*,*ω*_ > 0), and computed the proportion of above-threshold components shared between saccade and replay conditions. Whereas saccade and motion-only replay shared a considerable amount of 65.7% (*SD* = 23.3) of components, for the motion+suppression replay this overlap was significantly smaller, amounting to only 24.7% (*SD* = 24.4; *t* (5) = 4.54, *p* = .006). This difference was not unexpected, as simulated saccadic suppression by definition reduced the scenes’ power in the low-SF domain, rendering this theoretically and empirically extremely relevant source of information almost unresolvable. That suppression of low-SF information continued to be relevant beyond saccade offset can be appreciated in the rightmost panels of Figure 8e. Taken together, the description of smear in the SF-orientation by means of reverse regression led to highly similar results in saccade and replay experiments, provided that no suppression was applied. This result suggests that largely the same information was used to solve the scene-matching task in the two viewing modes. They also prove the point that simulated saccadic suppression indeed impaired those SF bands that were most crucial, which ultimately explains the heavy decrease in task performance in the combined motion and suppression replay.

## Discussion

Intra-saccadic smear is a ubiquitous consequence of the active perceptual exploration of the world, as it is necessarily induced every single time we make a saccade. The fascinating phenomenon that smear is yet not part of conscious visual experience does not preclude that it can be processed and perceived in principle: Building on the pioneer work of Campbell and Wurtz (1978), we tachistoscopically displayed scenes of natural environments while the eyes were in mid-flight to uncover the natural motion smear that – despite its physical presence – is otherwise omitted from perception.

### Intra-saccadic temporal dynamics of smear

In three separate, pre-registered experiments we mapped the temporal dynamics of motion smear throughout the course of the saccade, using both objective and subjective perceptual tasks. In a fourth pre-registered experiment, we simulated the visual consequences of saccades during fixation, to establish a baseline for potential extra-retinal influences on perceptual judgments. In both tasks, as well as both viewing conditions, perception of smear clearly followed an intra-saccadic time course. This suggests that saccadic omission of motion smear is not only a consequence of preand post-saccadic masking by largely stable, high-intensity retinal images present before and after saccades (Breitmeyer & Ganz, 1976; Castet, 2010; E. Matin, 1974; Volkmann, 1986), but that the same mechanism that generates smear during saccades could also contribute to its reduction. Put differently, masking of smear does not strictly start right after saccade offset – as Figure 2 in Campbell and Wurtz (1978) seemed to suggest – but is already taking place during the deceleration phase of saccades. In our experiments, deceleration phases were clearly longer than acceleration phases, a common finding for larger saccades (Van Opstal & Van Gisbergen, 1987). Only because saccadic velocity profiles are by nature continuous does not entail that the perceptual transition from saccade to fixation is also a smooth function. By incremental removal of variance in the saccade-duration distribution we however determined that perception of smear indeed followed a continuous time course. Interestingly, time courses of large-field stimuli were considerably steeper than those of Gaussian-window stimuli. This result matches real-world observations, as motion streaks, or even phantom arrays (Hershberger & Jordan, 1998), induced by small light sources in dark environments can remain conspicuous over longer periods after the end of the saccade (Jordan & Hershberger, 1994; E. Matin et al., 1972). The finding that larger Gaussian windows, which determined the width of the streak, facilitated the speed by which smear was alleviated around saccade offset can be considered evidence for the hypothesis that post-saccadic masking should be more efficient when more transient channels are stimulated around the end of the visual shift (Breitmeyer & Ganz, 1976; E. Matin, 1974). That is, when no strong mask is present when the saccade lands (for a similar argument, see Ibbotson & Cloherty, 2009), intra-saccadic signals can be seen for the duration of visual persistence. For instance, even very complex intra-saccadic retinal patterns can be reliably discriminated in sparse and dimly lit environments (Schweitzer, Watson, Watson, & Rolfs, 2019). Owing to the rich fullfield visual stimulation typically encountered in natural environments, a single RF that may have responded to an intra-saccadic transient will most definitely also respond to post-saccadic input. Even if only due to temporal recency, responses to post-saccadic input will thus inevitably outweigh responses to brief intra-saccadic signals.

### Model perspectives on the generation and reduction of smear

The idea that the temporal dynamics of smear are closely linked to the underlying saccade dynamics is supported by the two types of models presented in this paper, which viewed the process of reducing, or masking intrasaccadic smear from two alternative perspectives.

On the one hand, the process could be viewed as a direct consequence of the human contrast sensitivity profile. As retinal velocity decreases in the second half of the saccade, spectral power in the resolvable TF range gradually increases and thereby restores responsiveness to increasingly higher SFs. As a result, power at low SFs and orientations parallel to the direction of the saccade, which remain comparably high during saccades and therefore determine the appearance of smear, gradually dissipates. The same process restores access to mid- and high-SF content, such as fine spatial detail and edge information that can only be resolved during fixation (Martinez-Conde, Macknik, & Hubel, 2004; Rucci, Ahissar, & Burr, 2018). When this coarse-to-fine transition is concluded, smear is effectively eliminated, or masked.

On the other hand, the gradual reduction of smear can also be shown without taking human contrast sensitivity into account. In this case, the process would be a mere consequence of the saccade’s trajectory and the temporal dynamics of visual processing. We have shown that smear can be produced by simply averaging subsequently sampled inputs along a 2D trajectory, that is, the idea of a motion filter. Note that this model of smear shares the commonality with the above model that its output is dominated by orientations parallel to the defined trajectory. As eye velocity now decreases, subsequent inputs gradually become more similar to each other until finally, when the eye is approximately stable, the same inputs are being registered, ultimately overweighting the highly variable inputs received during the saccade. When this process is complete, smear is masked. For this model to work given an infinite time interval, it needs, but also benefits from the additional assumption of decaying activity, which is undoubtedly present in the early visual system (Albrecht, Geisler, Frazor, & Crane, 2002; Frazor, Albrecht, Geisler, & Crane, 2004; Reich, Mechler, & Victor, 2001).

Even though spatiotemporal and trajectory-dependent models are quite different, they are well compatible and could be combined. Trajectory-dependent motion-filter kernels were implemented to reproduce the saccade’s trajectory as a line and therefore still include a high-SF spectrum that would not remain resolvable during saccades, even if orientations were parallel to the saccade’s direction. Motion filters could thus be extended to incorporate the spatiotemporal processing characteristics of the visual system.

### Constraints of the proposed models

What both models have in common is that they are visual models and do not consider any extra-retinal factors that may have an influence on processing and perception of smear. First, they do not consider proactive modulations of neuronal responses in visual and motion-related areas that are specifically related to the motor command, and not the retinal shift (Bremmer, Kubischik, Hoffmann, & Krekelberg, 2009; Denagamage et al., 2023; Ibbotson, Crowder, Cloherty, Price, & Mustari, 2008; Sylvester, Haynes, & Rees, 2005; Thiele, Henning, Kubischik, & Hoffmann, 2002). Second, the models do not account for processes of visual localization. Transient stimuli presented during saccades are consistently localized and perceived in space, although this localization is prone to systematic errors (e.g., L. Matin & Pearce, 1965; Sperling & Speelman, 1965) – a phenomenon modeled assuming a damped representation of eye position (Dassonville, Schlag, & Schlag-Rey, 1992; Mateeff, 1978; Morris, Kubischik, Hoffmann, Krekelberg, & Bremmer, 2012; Pola, 2004). We indeed casually observed that smeared scenes appeared to be spatially shifted in the direction of the saccades, which is precisely what these models would predict. Although it would be an intriguing question to study to what extent eye position signals influence the generation and perception of smear, for reasons of parsimony we chose to treat smear as a purely retinotopic phenomenon. Third, we did not consider the question of motion perception. Subjectively, there is no feeling of a moving world during saccades, as opposed to saccade-like large-field motion presented on a screen or passive eye movements, for instance, when gently tapping or pressing the eyeball (e.g., Ilg, Bridgeman, & Hoffmann, 1989). Explaining this phenomenon of visual stability goes beyond the scope of our experiments and models, as it relates to the more general question of how the system detects and compensates for self-motion (Bridgeman, Van der Heijden, & Velichkovsky, 1994; MacKay, 1973). It should however be clear that, although sensitivity to motion is reduced during saccades due to suppression along the magnocellular and inferior pulvinar pathways (for a review, see Wurtz, 2008), perceiving motion during saccades is well possible, provided that stimuli are presented in a TF range suitable for motion perception (Castet et al., 2002; Castet & Masson, 2000; Nicolas et al., 2021). In our experiments, smeared scenes did convey a sense of motion on some occasions, but since we did not systematically survey these percepts, our models also did not include any dedicated detectors of motion (Adelson & Bergen, 1985; Neri, 2014; Reichardt, 1987) or motion streaks (Geisler, 1999), let alone their saccade-related modulations. Even though our models did not incorporate these important extra-retinal influences, they still did surprisingly well in reproducing the perceived content and extent of smear, which suggests that, in principle, the dynamics of perceived smear due to high-speed image translations across the retina could be a purely visual process.

### ‘Saccadic’ omission during fixation

Assuming that retinal dynamics alone are sufficient to understand the generation and reduction of intra-saccadic smear, the proposed models should be directly applicable to cases where image shifts occurring during saccades are simulated during fixation. Indeed, preand post“saccadic” masks were shown to be effective during fixation, for instance, in reducing motion amplitude (Duyck et al., 2018) or hindering target detection (Brooks, Impelman, & Lum, 1981; Chekaluk & Llewellyn, 1990), suggesting that the underlying mechanism may not be saccade-specific. We directly tested this hypothesis using a 1440fps projector capable of reproducing saccade dynamics and simulated saccadic suppression with high temporal fidelity. Replaying the visual consequences of saccades made in Experiments 1–2 and subsequently comparing results from saccade and replay experiments allowed the quantification of visual and non-visual contributions to the processing and perception of smear.

The striking similarity between saccade and (motion-only) replay experiments, as evident from the time courses and SF-orientation weight maps, provides strong evidence the dynamics of smear perception are indeed predominantly determined by the dynamics of saccade-induced visual input, making a strong case for the idea that “vision with the rapidly moving eye […] does not differ essentially from vision with the resting eye […] given only the same retinal stimulation” (Woodworth, as cited by E. Matin, 1974, p. 902). This statement can be misleading, as it was in fact not possible – or is in principle not possible – to achieve the exact same retinal stimulation across viewing conditions. For instance, there are intrinsic differences between saccade and fixation, ranging from inertial forces acting upon retinal photoreceptors (Richards, 1968) and the crystalline lens (Deubel & Bridgeman, 1995b), over neural modulations due to the motor efference (Bremmer et al., 2009; Denagamage et al., 2023; Ibbotson et al., 2008), to attention shifts and remapping (Cavanagh, Hunt, Afraz, & Rolfs, 2010; Rolfs, 2015). Furthermore, there are technical limitations of the experimental setup used for replay, such as static screen borders and the resulting partial occlusions of scenes, the reduced screen resolution and loss of color information, as well as measurement noise and artifacts introduced by eye tracking systems. Despite these limitations, smear generation and omission during active and simulated saccades were nearly identical. In the present study specifically, it may have been related to the (rather uncommon) use of natural scenes – large-field stimuli of considerable complexity that human observers have plenty of experience with – that saccades and replayed saccades were (also phenomenologically) so highly comparable. From these findings it seems fair to conclude that eye-movement-induced retinal dynamics, and especially the presence of a stable post-movement retinal image, can be considered the primary and sufficient criteria for the omission of motion smear during saccades.

### The role of saccadic suppression

The elephant in the room up to this point is the role of saccadic suppression. There is a large body of literature suggesting that this SF-specific reduction of contrast sensitivity around saccades is a phenomenon of extraretinal origin, triggered by the saccadic efference copy (e.g., Burr et al., 1994; Diamond et al., 2000; Riggs & Manning, 1982; Ross et al., 2001a; Sylvester et al., 2005). Replays that merely reproduce retinal motion trajectories during fixation would thus neglect that dedicated brain mechanisms might be in place to proactively dampen sensitivity to saccade-induced motion and smear, “precisely to avoid the disturbing consequences of saccadic image motion which would follow if it were left intact” (Burr & Ross, 1982, p. 483). In Experiment 4, therefore, we decided to simulate not only saccade-induced image motion but also saccadic suppression itself. Our reasoning was that, since active saccadic suppression – especially strong in presence of patterned scenes (Brooks & Fuchs, 1975; Diamond et al., 2000; Mitrani, Yakimoff, & Mateeff, 1973) – had contributed to the results in Experiments 1–3, a replay also taking into account such suppression would more closely approximate the saccadic scenario. It is thus puzzling that the opposite was the case: While motion-only replays produced results highly similar to those produced by saccades, combined motion and suppression caused literal gray-outs which rendered smear indiscernible, impairing task performance. These gray-outs are the exact perceptual consequence that should be expected from the combined activity of two processes: On the one hand, as shown by reverse-regression and modeling results, smearing degrades mid-to-high SFs (except those with orientations parallel to saccade direction). On the other hand, suppression takes care of the low SFs that can remain resolvable even at saccadic velocities (Burr & Ross, 1982). The problem is that such complete gray-outs were rarely experienced during our saccade experiments (see also Campbell & Wurtz, 1978), and were also not reflected in our results.

Why did adding the factor of saccadic suppression not contribute to creating a more veridical replay? From a technical standpoint, it could be argued that the strength of suppression was overestimated by the descriptive model applied in the replay. Whereas all data was adapted from the literature without further assumptions, it is indeed possible that the circumstances under which suppression was originally measured did not apply to our experimental context. For instance, Burr (1981) instructed almost twice the saccade amplitudes used in our study. Saccade amplitudes are known to modulate the amount of suppression, even though the size of this modulation was found to be 0.2 log units at most, when comparing 32-deg and 16-deg saccades (Stevenson, Volkmann, Kelly, & Riggs, 1986). Moreover, the data taken from Burr (1981) was measured at a higher background luminance than what could be presented in our studies. Indeed, saccadic-suppression magnitude scales with background luminance, however, dependence on background luminance is equally valid for suppression induced by simulated saccades (Brooks & Fuchs, 1975) and should therefore also affect replay conditions. A likely explanation, however, is that – in our paradigm specifically – saccadic suppression was simply not relevant. To elaborate, perceptual suppression was found to be virtually absent in total darkness (Mitrani & Yakimoff, 1971; Richards, 1969).Put differently, saccadic suppression might have been present but not of sufficient strength to differentially affect perceptual performance in saccade and replay conditions. In darkness, the difference between their respective thresholds amounts to 0.1 log units at most (Brooks & Fuchs, 1975), in which case the effect of smearing overrules the effect of suppression (Mitrani & Yakimoff, 1971). The simulated suppression we introduced in the replay, however, was significantly stronger than saccadic suppression measured in such low-light conditions, in which case suppression could overrule smearing.

From a more conceptual standpoint, an explanation for this puzzling effect of simulated suppression could be that the reduction of contrast sensitivity attributed to extraretinal saccadic suppression is confounded by, or even a consequence of visual processes. Such passive accounts of suppression have been discussed thoroughly before, especially in the face of the various visual factors that determine suppression magnitude (for excellent reviews, see Castet, 2010; E. Matin, 1974; Volkmann, 1986). Recently, Idrees et al. (2020) found that saccadic suppression was already present in the in ex vivo retinae as a consequence of image motion and that SF-selectivity of suppression was modulated by the SF-content of the visual periphery, both during saccades and simulated saccades. Applying this argument to the experiments at hand, saccadic suppression – due to its retinal origin – has already occurred to its full extent when the large-field motion of the visual scene was presented. Therefore, it did not matter whether visual stimulation was elicited by real or simulated saccades, and, for the same reason, any additional extraretinal reduction of contrast sensitivity would only impair performance beyond the expectable level. While our replay experiments are clearly unfit to provide insights in the question about the origin of active saccadic suppression, they do provide one additional major piece of evidence: Omission occurred not only independently of active saccades but also in the absence of (very powerful) contrast suppression. Thus, saccadic suppression, at least in its classic active formulation, explains neither our results, nor the overarching phenomenon of saccadic omission.

### Active contributions to saccadic omission

Finally, passive views are obvious antipodes to active views, which invoke efference-based prediction processes to explain saccadic omission. The idea is that, thanks to motor information and repeated experience, the visual system has an accurate representation of the sensory consequences of its own saccades, allowing it to omit expected and render visible unexpected peri-saccadic information. Indeed, there is good evidence that habituation to saccade-contingent stimulation can lead to the upregulation of detection thresholds for the same or similar stimulation (Pomè, Schlichting, Fritz, & Zimmermann, 2024; Zimmermann, 2020), even though classic contrast suppression seems to be an exception to this finding (Zimmermann & Lange, 2022). Adopting this active view, it could be argued that, by removing the (normally present and thus expected) post-saccadic view of the scene, a prediction error is introduced that renders saccade-induced visual input visible. In the presence of a post-saccadic scene, however, no such prediction violation occurs and saccade-induced input is omitted. This view is arguably less parsimonious and remains unspecific about several aspects. For one, it is unclear how a prediction about the visual consequences of the scene’s rapid shift over the retina should be generated when the stimulus is only revealed after the saccade has been initiated. Certainly, the contingency that a scene would be presented upon saccade execution could be learned throughout the course of the experiment. The scene identity, however, or rather its smeared appearance, remained unpredictable to observers on each single trial. Arguably, the notion of ‘prediction’ could be a generic one – that is, the system could expect average natural-scene statistics. Whether such a prediction would be sufficient to perform omission, given the wide range of natural scenes we presented and the systematic differences between scene categories we observed, is unclear. The active view also tacitly assumes that, in order to effectively omit expected visual input induced by saccades, the system has a time-resolved representation of the temporal dynamics by which visual information is typically modulated throughout the saccade described by the motor command. While it is one of the major goals of this paper to understand these spatiotemporal dynamics, and whether the visual system could in principle make use of some of them, it still seems rather diffi-cult and computationally expensive to achieve such realtime correction, especially in the light of the great diversity of visual environments and the wide range of visual transients (potentially with varying latencies) they induce when in motion. Finally, it is not entirely clear how active omission could function in the absence of an efference copy, that is, when no saccades are made and still remarkably similar omission of smear occurs, as shown in our replay experiment, as well as other simulated saccade studies (e.g., Chekaluk & Llewellyn, 1990; Duyck et al., 2018). On the one hand, it may be that, in such special cases, active omission is simply not at play and that other (e.g., passive) mechanisms are in place, which produce virtually the same effects. On the other hand, the visual system may in principle be capable of performing active omission by postdiction. In this slightly adapted version of active omission, the system would assert – regardless of actual saccade execution – whether a certain pattern of visual stimulation is compatible with the stimulation introduced by a saccade and, if so, omit it from perception. This view would have the advantage that no real-time prediction error would have to be computed and visual information could be integrated over a somewhat longer interval. Preliminary evidence consistent with this view was provided by Rolfs, Schweitzer, Castet, Watson, and Ohl (2023) who showed that thresholds for detecting high-speed target motion during fixation could be predicted by the main-sequence law of saccade kinematics. That said, active efference-based prediction (or postdiction, see Masselink & Lappe, 2021) undoubtedly plays a major role in trans-saccadic vision, motor learning, and visual stability (Wurtz, 2018), and to see these principles extended to the omission of saccade-induced sensory input would certainly be intriguing. The question whether they work in conjunction with, or should even be preferred over passive mechanisms, which require little more than early visual processing, is a matter of future research.

### Experience of saccade-induced smear

The spatiotemporal models proposed in this paper suggested that, as a consequence of human contrast sensitivity and the velocities of the retinal shift, only a small subset of SFs and orientations present in natural scenes could actually be resolved during saccades and simulated saccades – a notion well supported empirically by the respective reverse-regression analyses.

First, if oriented orthogonal to their motion direction, contrast sensitivity to (very) low SFs increases when they are presented at saccadic velocities (Burr & Ross, 1982), whereas higher SFs presented at those velocities produce TFs around or above their fusion threshold and are thereby rendered ‘invisibile’ (Castet & Masson, 2000; García-Pérez & Peli, 2001, 2011). Note that for this reason and in order to reduce the impact of retinal velocity on measurements of contrast sensitivity, many saccadic-suppression studies in the past presented horizontal gratings during horizontal saccades (e.g., Burr et al., 1982, 1994; Niemeyer et al., 2022; Volkmann et al., 1978).

Second, orientations parallel to the direction of the retinal shift move at considerably lower velocities, which is why, for these orientations, it is possible to resolve much higher SFs. This was predicted by an extension of the contrast sensitivity function by Kelly (1979) and confirmed by reverse-regression results from Experiments 1, 2, and 4: Scenes that contained higher power in orientations parallel to the saccade trajectory were more likely to be correctly identified, as these scenes’ defining features were more likely to remain intact during presentation. The relevance of these parallel orientations share interesting similarities to the motion-streak system. Motion-streak detectors integrate orientations orthogonal to direction-selective motion RFs, that is, orientations parallel to the direction of detectable motion (Geisler, 1999). This particular configuration could allow a motion-streak detector to encode fast motion of a dot stimulus as an orientation (Alais, Apthorp, Karmann, & Cass, 2011; Apthorp, Cass, & Alais, 2010, 2011; Apthorp et al., 2013; Geisler et al., 2001; Jancke, 2000). While motion streaks have been studied using stimulus velocities well below saccadic velocities, recent studies showed that intra-saccadic motion streaks, too, were enhanced if inducing stimuli contained orientations parallel to their retinotopic motion trajectory (Schweitzer & Rolfs, 2020b, 2021). Here we demonstrated that the predominance of parallel orientations also holds for natural scenes.

Third, another prominent feature of smear in natural scenes was its color content. Throughout all experiments we found that color had a consistent additive effect on all dependent variables, that is, it did not alter observed time courses. The effect of color is not surprising in the case of scene-matching performance, as this additional feature, shown to be somewhat less impacted during saccades (Braun et al., 2017; Burr et al., 1994; Knöll et al., 2011), would provide an additional cue to solve the task. In the case of filter adjustment, however, it was more surprising to see that the extent of perceived smear was reduced in colored images. In principle, it is possible that mechanisms of color processing interacted with the processing of smear. Indeed, color information may be integrated along a motion trajectory (Nishida, Watanabe, Kuriki, & Tokimoto, 2007), although it is unclear whether this finding could hold for saccadic speeds. Temporal sensitivity profiles for color were also shown to be more low-pass than those for luminance (Kelly, 1975), suggesting that, when combined, sensitivity could be improved at low TFs. Alternatively, it may seem more likely that the amount of perceived smear was overestimated in reference to the amount of smear that could have induced given the amount of the retinal shift. That this overestimation is plausible was demonstrated by the trajectory-dependent motion-filter model. While the source of this overestimation is unclear, we believe it is either a limitation of the experimental task, which only allowed observers to respond by adjusting the parameters of perfectly linear motion, or of perceptual origin. Specifically, linear motion filters leave high-SF information largely intact, while we have shown that their power is drastically reduced. Participants might have over-adjusted motion filters to better account for this inherent difference. Thus, the reason why perceived smear was reduced in color images may be simply that color information reduced the perceptual bias of overestimating smear. Indeed, color and luminance edges were shown to be relatively independent in natural scenes (Hansen & Gegenfurtner, 2009) and could therefore have provided additional structure to the scene, making it easier for observers to veridically reproduce the smeared scene. This benefit has been reported for static scenes presented for durations as short as the ones applied in our experiments (Gegenfurtner & Rieger, 2000). In sum, our results suggest that saccade-induced smear could be conceptualized as an orientation-SF bandpass filter that restricts processing to low-SF content but allows higher SFs around orientations close to each saccade’s direction, as well as color, to pass. The extent of this filter was however not only related to the instantaneous velocity of the retinal shift – as the function by Kelly (1979) would suggest. It was also – as the trajectory-dependent motion-filter model would suggest – critically dependent on the amplitude of the shift, which increased monotonically with presentation duration in our paradigm.

### Potential functional relevance of saccade-induced smear

Given the efficiency and parsimony of preand postsaccadic masking, saccade-induced motion smear will, at least in natural environments, rarely impact or impair our subjective perception of the world. It has been demonstrated that, even during fixation, static endpoints of as little as 50 ms of duration were capable of rendering the shape of a rapidly moving object indiscriminable (Wexler & Cavanagh, 2019). Similarly, our models, as well as results by Campbell and Wurtz (1978), suggest a virtual absence of smear as early as 30 ms after saccade offset. It may be interesting to note that due to the presence of post-saccadic oscillations, which are especially prominent in the lens, the retinal image is still be moving even after the eyeball has finished its rotation (Tabernero & Artal, 2014). Such movement has been shown to have perceptual consequences – even though they are much smaller than the size of the measured PSO in the lens (Deubel & Bridgeman, 1995a, 1995b) – and may add to the explanation why large-field smear was still perceived reliably even up to 20 ms after saccade offset (Campbell & Wurtz, 1978).

In the light of these findings one may ask why one should even care what saccade-induced smear looks like? For one, saccade-induced smear is clearly part of the visual transients that facilitate a coarse-to-fine processing strategy of the novel post-saccadic input (Rucci et al., 2018). Assuming a delayed temporal response, saccades enhance low-SF input during early fixation, whereas fixational eye movements enhance high-SF input during late fixation (Boi, Poletti, Victor, & Rucci, 2017). The bandwidth of this enhancement depends on the amplitude of each saccade (Mostofi et al., 2020) and may be particularly relevant in natural scenes where power is inherently high in the low-SF domain. Our results suggest that, in addition to a redistribution of power in SF space, there is a redistribution of power in orientation space depending on the direction of each saccade. Curiously, both distributions of saccade directions and power in natural scenes are strongly biased towards cardinal orientations (Rolfs & Schweitzer, 2022) – a similarity that an oculomotor strategy could in principle exploit. Furthermore, knowing what type of input could be expected from saccades could in principle help the visual system to omit this particular input from conscious perception. Recent evidence suggests that perisaccadic visual input is not discarded but actively monitored, for instance, in order to adjust the profile of saccadic suppression (Scholes, McGraw, & Roach, 2021; Zimmermann, 2020) or to inform gaze correction (Schweitzer & Rolfs, 2021). These findings point towards a potential relevance of saccade-induced visual input, but it should be noted that virtually all studies in this domain use detectable transients during saccades to assess visual function. While these manipulations are necessary, which is clearly also the case in this study, they are not ecologically valid: It would in fact rarely ever happen that a single target stimulus or even the entire visual scene shifted in a direction perpendicular to the direction of the saccade – and, if so, such orthogonal steps would be readily noticed, and not omitted (Wexler & Collins, 2014). If the visual system drew upon sensorimotor contingencies to perform saccadic omission (e.g., Pomè et al., 2024; Zimmermann, 2020) then it is likely they would be established in natural environments. Rather than dealing with transient stimuli like flashes or abrupt displacements, processing should then be optimized for continuous stimuli (for similar arguments, see Schlag & Schlag-Rey, 1995; Teichert, Klingenhoefer, Wachtler, & Bremmer, 2010) that reflect inherently continuous retinal shifts induced by saccades across largely static scenes. From this point of view one could argue that little is actually known about saccadic omission under non-laboratory circumstances, that is, with both natural and continuous stimuli. Thus, studying the features and temporal dynamics of the visual consequences that should occur during natural vision may be one first step to understand how this impressive feat is performed.

## Conclusion

How the visual system gets rid of motion smear is a long-standing question. Motion smear induced at moderate velocities, such as the one expected when taking a photo of a moving object at low shutter speeds, is greatly reduced at longer presentation durations as a consequence of motion-selective mechanisms (Burr, 1980). In contrast, motion smear induced by saccades – movements easily a tenfold faster than typical object velocities found in natural moving images (Dong & Atick, 1995) – can be readily perceived, so that, in order to reduce it, different sets of mechanisms must come into play. Resolving the temporal dynamics of perceived motion smear throughout and after saccades, our results show that the extent of smear follows a continuous, intra-saccadic time course closely linked to the inducing saccade’s profile. Crucially, not only generation and (post-saccadic) masking of smear but also the subjective appearance of smear could be modeled by the same process, which needed only few ingredients to work: The saccade’s trajectory as well as the temporal properties and contrast sensitivity profile of the visual system. These same ingredients were capable of producing virtually identical omission when no real saccade but only saccade-like motion was present. These results point towards an intriguing perspective: Omission of saccade-induced smear could be understood as a consequence of the interplay between saccade-induced retinal dynamics and early visual processing. This parsimonious mechanism would rely only on the processing hardware of the early visual system to eliminate smear. As such, it would greatly reduce the computational load on the sensorimotor system when combining rapidly changing visual information with motor and proprioceptive signals to achieve visual stability.

## Methods

### Apparatus

#### Experiments 1–3

All experiments were conducted in a sound-attenuated, dark room. Stimuli were presented on a gamma-linearized, 52 cm wide, 16:9 VIEWPixx 3D monitor (VPixx Technologies, Saint-Bruno, QC, Canada) with a resolution of 1920 x 1080 pixels at a refresh rate of 120 Hz. Participants were seated at a viewing distance of 57 cm with their head supported by a chin-rest. Eye movements of the dominant eye were recorded with Eyelink 1000 desktop system (SR Research, Osgoode, ON, Canada) at a sampling rate of 1000 Hz with no link filter and normal data filter. Responses were collected with a standard USenglish keyboard. All experimental code was implemented in MATLAB 2016b (Mathworks, Natick, MA, USA), using Psychtoolbox (Kleiner et al., 2007) and the Eyelink toolbox (Cornelisen, Peters, & Palmer, 2002), and ran on a workstation with a Ubuntu 18.04 operating system. For experiments 1 and 2, a NVIDIA GeForce GT740 graphics card was used, whereas for experiment 3 – to accelerate convolutions with motion filters – a Geforce RTX2060 was installed.

#### Experiment 4

Stimuli were displayed onto a 16:9 (220 cm width) video-projection screen, mounted 270 cm in front of the participant, using a by default gamma-linearized PROPixx DLP projector (VPixx Technologies, Saint-Bruno, QC, Canada) running at 1440 Hz vertical refresh rate and a resolution of 960 x 540 pixels and a deterministic latency equal to 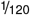 seconds. As this projector changes all screen pixels at the exact same time, presentations were directly comparably to the previously employed tachistoscopic presentation. Again, eye movements of participants’ dominant eye were recorded at 1000 Hz (normal link and data filter settings), this time using an Eyelink 1000+ desktop system (SR Research, Osgoode, ON, Canada), running MATLAB 2022b on Ubuntu 20.04.

### Tachistoscopic display

Whereas the timing of the PROPixx projector has been evaluated (Schweitzer & Rolfs, 2020a), the timing of the VIEWPixx monitor’s tachistoscopic mode, possibly due to its rare use in scientific research, has not. To this end, we used a LM03 lightmeter (Cambridge Research Systems Ltd., Rochester, UK) recording at a sampling rate of 2000 Hz, which was similarly used in previous screen evaluations (e.g., Shi, 2017). We displayed on-off-flicker at four different on-durations (8.33, 16.67, 25.0, and 33.33 ms, i.e., the presentation durations in Experiment 1), separated by a constant off-duration of 8.33 ms, which resulted in a total of 1200 trials for each on-duration. Results revealed that within the temporal resolution of the lightmeter, that is, 0.5 ms measured on-durations were precisely as specified by the schedule (Figure 1b). To measure the onset latency of the screen following the tachistoscopic trigger, we capitalized on the fact that the LM03 lightmeter uses two separate photosensors and implemented the presentation schedule in a way to turn off a reference LED upon the trigger onset. We found that the on-set of the screen was at the same time as, or even earlier than the LED offset (*M*_*offset*_ = − 0.16 *ms, SD* = 0.24 *ms*), suggesting that latencies of tachistoscopic presentations were largely negligible. Minimal code to drive the LM03 lightmeter can be found at https://github.com/richardschweitzer/LM03_lightmeter. The scripts used to evaluate the tachistoscopic display are provided on the OSF project page https://osf.io/bf246/. To synchronize visual presentation onsets and eye tracking data, messages were sent to the Eyelink host computer via the TCP link as soon as presentation schedules were initiated.

### Participants

20 observers were tested in each experiment. This sample size was determined based on power contours computed with an available online tool designed to plot the relationship between numbers of participants and trials (Baker et al., 2021). We retrospectively estimated the feasibility of this procedure after completing data collection of Experiment 1: Using 45 trials per experimental cell and observer (see Experimental design), resulted in statistical power well above 80%, given *α* = 0.05, estimated *σ*_*b*_ = 0.094 and *σ*_*w*_ = 0.049, and assuming a difference of means amounting to 0.066 (i.e., the mean difference in the proportion of correct responses for display durations of 8.33 and 33.33 ms in Experiment 1; see Figure 3b).

Prior to inclusion in the study, all gave written informed consent. Visual acuity test established normal or corrected-to-normal vision (i.e., 20/20 ft. acuity in the Snellen test) and ocular dominance was tested with a variant of the Porta test. All experiments were preregistered and conducted in agreement with the latest version of the Declaration of Helsinki (2013) and approved by the Ethics board of the Department of Psychology at Humboldt-Universität zu Berlin (protocol number 2018-36, “Mechanismen visueller Stabilität”). As remuneration participants received 8 Euros per hour, plus 2 Euros for every 15 minutes of overtime.

#### Experiment 1

14 female and 6 male participants in the age range of 19 to 33 years (M = 24.36, SD = 4.6) took part in the study. 16 out of 20 participants had right ocular dominance and all were right-handed. Seven wore glasses and one participant wore contact lenses. In accordance with preregistered exclusion criteria, three participants were excluded and then replaced throughout data collection due to insufficient quality of eye tracking data. Pre-registration and data can be found at https://osf.io/bf246/.

#### Experiment 2

11 female and 9 male participants in the age range of 21 to 36 years (M = 26.35, SD = 4.45) participated. 15 had right ocular dominance and 19 were right-handed. Nine participants wore glasses. One participant had to be excluded and replaced due to poor eye tracking. Preregistration and data can be found at https://osf.io/ue6cd.

#### Experiment 3

13 female and 7 male participants in the age range of 22 to 35 years (M = 27.25, SD = 3.24) participated in two sessions, each with a duration of one hour, on separate days. Again, 15 had right ocular dominance and 19 were right-handed. Eight participants wore glasses and three contact lenses. Pre-registration and data can be found at https://osf.io/ravbc/.

#### Experiment 4

20 participants (15 female, 4 male, 1 diverse) in the age range of 21 to 63 years (M = 30.65, SD = 10.08) participated in two sessions, again with a duration of one hour and on separate days. 17 had right ocular dominance, 19 were right-handed. Seven participants wore glasses, two contact lenses. Pre-registration and data can be found at https://osf.io/56mqg/.

### Task procedure in saccade experiments

Each trial started with the display of the fixation marker on the left side of screen at 8 degrees of visual angle (dva) horizontal eccentricity relative to screen center, as well as the saccade target marker on the right side of screen at 8 dva horizontal eccentricity, on a black background (see Stimuli). This procedure resulted in the instruction of a perfectly horizontal saccade amplitude of 16 dva in Experiments 1 and 2. In Experiment 3, in order to make motion-filter angle adjustments (see below) more task-relevant, saccade directions were uniformly sampled from +22 to -22 degrees relative to horizontal. To pass fixation control, 450 ms of successful fixation within a circular boundary with a radius of 2 dva around the center of the fixation marker were needed. Trials could be aborted in case of 2 seconds of not fixating within the boundary or more than 20 re-fixations after repeatedly breaking fixation. As both of these cases suggest poor eye-tracking accuracy or precision, they automatically triggered calibrations. Upon successful fixation, the backlight of the monitor was turned off, constituting the cue to make a saccade to the previously presented saccade target. As soon as the backlight was turned off, the image of the first scene was drawn to the screen to achieve an earlier and more reliable gaze-contingent display onset. Saccades were detected online using the algorithm proposed by Schweitzer and Rolfs (2020a) using the parameters *λ* = 15, *k* = 2, and *θ* = 40. Online saccade detection triggered the tachistoscopic display schedule, that is, the backlight of the screen was turned on for the defined display duration (see Experimental design) and then turned back off, thereby presenting the first scene image (Figure 1a). Trials were aborted if no saccade was detected within a time window of 2.5 seconds after cue onset.

#### Experiments 1 and 2

The presentation of the second scene either the same or a (randomly selected) different scene was initiated after an inter-stimulus-interval (ISI) of 500 ms. This second scene was presented using the same display duration and color condition. Observers responded using RightArrow to indicate that they believed that the two scenes were the same and LeftArrow to indicate that they were different scenes. This response also triggers the reactivation of the monitor’s scanning backlight, presenting solely the initial saccade target marker.

#### Experiment 3

Following the ISI, the previously presented scene reappeared, prompting observers “to adjust a motion filter to make the image look as similar as possible to the one perceived during the saccade” (participant instructions). The filter was adjusted using ArrowRight and ArrowLeft to increase and decrease the length of the motion filter, thereby increasing and decreasing the amount of smear, whereas the keys ArrowUp and ArrowDown controlled the angle of the filter, adding upward and downward components to the blurred image. By default, the image appeared with no motion filter applied, so that observers always viewed a clean version of the scene. A video elucidating this task procedure can be found at https://osf.io/dj8xw/. To speed up convolutions and give observers the possibility to experience changes in the motion filter in near real-time, motion filters were run on the graphics card and on an image matrix downscaled to 50% of its initial resolution, which was restored to its initial dimension for presentation. Motion-filter angle could be adjusted at steps of 1 degree, whereas length could be adjusted at steps of 10 pixels, amounting to 0.27 dva. Observers had unlimited time to adjust the motion filter and pressed Enter once they were satisfied with the image.

#### Trial repeats

Post-trial text feedback, as well as a visual display of the saccade trajectory, was provided if gaze position did not reach a 5 dva circular boundary around the saccade target (‘Saccade did not reach the target area’), if two or more saccades were made to reach that boundary (‘Please make one saccade (you made X)’), or if a wrong response key was pressed (‘Wrong response key pressed’). Trials, in which any of these events were detected, no saccade was made, or fixation control was not passed, were repeated at the end of all regular trials. Each session will be divided in 8 blocks.

### Task procedure in fixation experiment

Experiment 4 was designed as a replay version of Experiments 1 and 2. Each trial started with the presentation of a fixation marker around screen center, displayed on a black background. After successful fixation control (identical to saccade experiments), the fixation marker was turned off and the replay sequence was initiated. Note that the projector’s lamps were not turned off, but instead all pixels were set to minimum-luminance black. This was done because the projector’s lamps, unlike the backlight of the VIEWPixx monitor used in Experiments 1–3 (see evaluation of the Tachistoscopic display), have been shown to only reach half of their maximum luminance after 13 ms and their full luminance after 70 ms (Duyck, Collins, & Wexler, 2021), rendering a backlight manipulation impractical. Around the time of display onset (as recorded in the original saccade trial) the scene was presented for the display duration while being shifted across the screen according to the previously registered saccade trajectory (see Simulated saccades). The second, post-“saccadic” presentation – again, either the same or a different visual scene – was initiated 500 ms after the offset of the first scene and presented centrally for the same display duration as the first scene. 250 ms after the second presentation, observers were allowed to respond using the same keys as in Experiments 1 and 2.

#### Trial repeats

Feedback was presented if initial fixation control was not passed (‘Please fixate’), if eye movements were made outside of a 3 dva circular boundary around the fixation target (‘Please do not make saccades’), if a wrong response key was pressed (‘Wrong response key pressed.’), or if a key press was registered before the presentation schedule, that is, the presentation of the first and second scene, was concluded (‘Please wait with your response until the trial ended!’). Any of the above events led to a repetition of the trial at the end of the session.

### Stimuli

Fixation markers were maximum-luminance, white squares 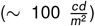 with black center of 0.4 dva width and height. Saccade target markers were identical but of twice the size of fixation markers. In Experiments 1– 3, background color was set to minimum-luminance black 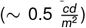 to minimize the luminance step that occurred when the monitor’s backlight was turned off. Note that these luminance measurements were conducted when the scanning backlight of the monitor was active. During tachistoscopic presentations, however, standard backlight mode is used, which increases overall luminance by a factor of 2.5, approximately. Luminance values in Experiment 4 were comparable, i.e., 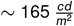 at maximum luminance.

Natural images were extracted from the SYNS database (Adams et al., 2016), that is, nine nonoverlapping single images per taken from each panoramic image in the database (https://syns.soton.ac.uk/). Single images were rescaled to a resolution of 1921 x 1281 pixels, using bicubic interpolation and thereby keeping their initial aspect ratio, and then converted to JPG images. During presentation the center of the image was aligned with the center of the screen, which cropped the image by 100 pixels on the upper and lower end to avoid a distortion of the image due to the slightly lower vertical screen resolution. In Experiment 4, where color channels were deliberately sacrificed and spatial resolution halved in order to achieve a refresh rate of 1440 fps, images were further downsampled to 960 x 540 pixels, so that they could be presented on the entire screen. Images were presented across the entire range of the monitor, thus covering approximately 49.0 x 28.7 dva in Experiments 1–3 and 44.3 x 25.8 dva in Experiment 4. All extracted scene images were deposited on OSF: https://osf.io/gxpuy/. Grayscale images were computed according to the weighted sum *I* = 0.299*r* + 0.587*g* + 0.114*b*, where *r*, *g, b* are the pixel values in red, green, and blue color channels of the original image.

As the SYNS database contained 90 panoramic images, extracting nine non-overlapping images resulted in nine sets of 90 images each, out of which eight (and in Experiment 4 all) were randomly selected for every single experimental session.

#### Experiments 1 and 2

Each set was shuffled and then randomly assigned to a color x duration sub-condition (2 x 4 conditions in total), for which the set was split into two halfs: One in which the first scene was presented with a matching second scene and one in which it was presented with a non-matching second scene. Each first scene of a set was therefore presented once per experiment, whereas each second scene was presented twice, that is, once with a matching and once with a non-matching first scenes.

#### Experiment 3

Each set was shuffled and then randomly assigned to a duration condition (8 conditions in total). For each set, a scene stimulus was randomly assigned a color x window sub-condition. Crucially, a window condition was added in Experiment 3 (see Experimental design). Scene images covered either the entire screen (full-field condition) or rendered as if they were seen through a Gaussian window (see example in Figure 1a). In this window condition, image matrices were multiplied with 2D Gaussian masks so that only a small, randomly chose portion of the scene remained visible. Gaussian windows had standard deviations of 0.5, 1, 1.5, 2, or 2.5 dva, which were randomly assigned in each trial. Window positions were also chosen randomly, following a uniform distribution in the range of ± 8.7 dva horizontal and ± 5.8 dva vertical eccentricity relative to image center.

#### Experiment 4

Procedures to create image sets followed those established in Experiments 1–2, with the exception that each of the nine image sets was associated with one display duration and was therefore presented twice, that is, once in each filter condition (see below). Note that the color manipulation present in previous experiments was dropped and all scene stimuli were presented in grayscale, as no color channels were available in the projection system’s 1440-fps mode.

### Simulated saccadic suppression

In Experiment 4 we introduced a simulated saccadic-suppression condition. To emulate a reduction of contrast sensitivity that affects specific SF bands and follows a time course determined by previous saccadic-suppression studies, we used the model developed in this manuscript (see Saccadic suppression for model details) to compute the suppression ratio for each SF contained in the image (Figure A4b), separately for each time point. To model the saccade onset-locked time course of saccadic suppression, we used the model parameters approximating contrast sensitivity measured with uniform backgrounds (Figure A4a), as those results should be largely uncontaminated by visual events, such as large-field motion (Diamond et al., 2000). Finally, power in each image’s SF domain was reduced (or enhanced) by dividing each complex value of the Fourier-transformed image by the corresponding suppression ratio. This procedure was performed for every frame of the replay sequence, thereby creating a time-resolved filter mimicking the SF-specific effects of saccadic suppression. Note that this procedure made sure that the average luminance of the image did not change by leaving the DC component of the image unchanged. An example video of a static pink-noise texture being filtered according to a full saccadic-suppression cycle can be found at https://osf.io/xh5qd.

#### Simulated saccades

To implement a replay capable of replicating the exact saccadic velocity profiles and timing of stimuli during fixation, saccades were randomly selected from valid trials collected in Experiments 1 and 2, excluding the outer 2.5% and 97.5% quantiles of saccade amplitude, duration, and latency distributions, to omit the most extreme motion trajectories from replay. Replay sequences started at the time of the recorded trial’s cue onset, thus replicating the saccade latency recorded in saccade trials, and ended at 110 ms after saccade onset – a value chosen to exceed both the longest saccade duration and the latest display offset after saccade offset. Display onset was triggered according to the timing of display onsets and offsets relative to the saccade onset measured in each saccade trial. To take into account possible systematic variations in replayed trajectories, replay sequences were also matched to display duration conditions according to the display duration in saccade trials, especially because short display durations could slightly shorten saccade trajectories (Figure 2c-e and Figure A3). For display durations longer than those tested in saccade experiments, saccade trajectories were randomly selected from trials collected in Experiment 2. To avoid repetitions in simulated-saccade trajectories, each saccade trial could only be sampled once for replay.

To match the projector’s high temporal resolution, the gaze signal was spline-interpolated and upsampled to 1440 Hz. Retinal trajectories were computed by subtracting gaze coordinates from the screen’s center coordinates and converting to degrees of visual angle to match the different specifications of the new setup. Owing to the fact that in Experiments 1 and 2 saccades were always made across the center of the screen, the full-screen scene would also always travel across screen center during replay, even though, due to the limitations of the screen in terms of visual angle, the scene was not always presented in its entirety throughout its motion trajectory – an unavoidable side-effect which was however barely noticed given the velocity and brevity of the scene’s motion. All saccade trajectories used for replay, as well as the scripts used to create them, can be found at https://osf.io/dynu7/.

### Experimental design

#### Experiments 1 and 2

One session per participant was conducted, containing 720 trials. Four tachistoscopic display durations were tested, that is, 8.3, 16.7, 25, and 33.3 ms in Experiment 1 and 33.3, 41.7, 50, and 58.3 ms in Experiment 2. Stimuli were either presented as color or grayscale images. First scenes (i.e., saccade-triggered presentations) and second scenes (i.e., scenes presented after the ISI) were either the same or different. Given this design, 45 trials were tested per experimental cell.

#### Experiment 3

Two sessions per participant were tested. Each session consisted of 352 trials, resulting in 704 trials in total. Eight display durations were applied, i.e., 8.3, 16.7, 25, 33.3, 41.7, 50, 58.3, and 66.7 ms, and, as in Experiments 1 and 2, color and grayscale scenes were shown. In addition, images could be presented as full-field images or as local spots described by a Gaussian window (see Stimuli). Consequently, observers completed 22 trials per experimental cell.

#### Experiment 4

Two sessions per participants were tested, each consisting of 810 trials. Nine equally spaced display durations ranging from 8.3 to 75 ms were used, thus – like in Experiments 1 and 2 – resulting in 90 trials (45 same and 45 different scene pairs) per duration. The additional factor of simulated saccadic suppression was introduced, so that each scene was presented twice, once in their original and once in their filtered form. All trials (in all experiments) were presented in a randomly interleaved fashion.

### Analyses

#### Pre-processing for saccade experiments

Eye-tracking data recorded at 1000 Hz was downsampled to 500 Hz and saccade detection was performed using the velocity-based Engbert-Kliegl algorithm (Engbert & Kliegl, 2003; Engbert & Mergenthaler, 2006) with *λ* = 5 and a minimum event duration of 3 samples. According to the processing pipeline described by Schweitzer and Rolfs (2022), we detected the onsets of post-saccadic oscillations (PSOs), based on which we then defined saccade duration. In case PSOs could not be detected, which may well happen when the saccade component containing the PSO is too short or too slow to be detected in the first place, we used the saccade offset determined by the Engbert-Kliegl algorithm. Saccade candidates that were no further apart than 5 samples were merged into one saccade. Within these merged saccades, PSO onsets were found using the method of direction-inversion (threshold: 70 degrees), and, if no inversion could be found, minimum velocity served as a criterion for PSOs (Schweitzer & Rolfs, 2022). PSOs were detected in 93.3% (range 34.4%–99.4%) of trials in Experiment 1, 89.8% (range 36.4%–99.8%) of trials in Experiment 2, and 94.8% (range 65.5%–99.8%) of trials in Experiment 3.

Trials had to pass four criteria to be included for analysis: (1) Fixation was passed and at least one saccade could be detected, (2) display onset of the first scene occurred during a saccade and display onset of the second scene occurred during fixation, (3) saccade candidates contained no missing or invalid samples, and (4) saccade metrics were compatible with an observer’s individual main-sequence relationship (Bahill, Clark, & Stark, 1975). This fourth criterion was necessary to remove saccades with extreme metrics, that is, unrealistically high peak velocities or overly large saccade amplitudes, while accounting for the atypical shapes of some observers’ velocity profiles (Figure A1). To approximate the individual main sequence of an observer, the amplitude-duration relationship was fitted using the square-root model

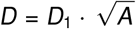

(Lebedev, Van Gelder, & Tsui, 1996) and the amplitude-peak velocity relationship was fitted by the exponential model

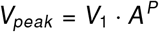

(Becker, 1989), where *A* denotes saccade amplitude. Both models were fitted in a robust manner, using an iterated reweighted least squares approach (Todorov & Filzmoser, 2009), to minimize the impact of outliers. Trials were excluded if saccades diverged more by more than 150 deg/s and 30 ms from the estimated main-sequence relation. Following this procedure, 94.2% (range: 77.7%– 99.6%) of trials in Experiment 1, 97.2% (range: 93.1%–99.6%) of trials in Experiment 2, and 96.4% (range: 79.6%–100.0%) of trials in Experiment 3 passed criteria 1–3. Of the remaining trials, 89.7% (range: 28.0%–98.6%) in Experiment 1, 94.0% (range: 74.0%–99.4%) in Experiment 2, and 88.6% (range: 42.6%–98.8%) in Experiment 3 also passed criterion 4, resulting in a total of 727 (range: 203–1231) trials in Experiment 1, 749 (range: 550–1078) trials in Experiment 2, and 676 (range: 313–868) trials per observer in Experiment 3.

#### Pre-processing for fixation experiment

As a first step, trials were excluded if participants had not passed initial fixation control, pressed a wrong response key or responded preemptively while stimulus presentation was still ongoing, if participants’ gaze exited the initial fixation region during stimulus presentation, or if presentation frames were dropped. This first step resulted in an exclusion of 2.6% (range: 0.2%–11.6%) of trials. Second, saccade detection was performed on eye-tracking data downsampled to 500 Hz, again using the Engbert-Kliegl algorithm with *λ* = 5 and a minimum event duration of 6 samples. Owing to the fact that the goal of saccade detection in Experiment 4 was merely to exclude trials in which saccades had occurred, neither PSO detection nor an analysis of saccade dynamics was performed. Specifically, trials were removed if there was any temporal overlap between saccades and presentations of the first or second scene, which occurred in 2.4% (range: 0.06%– 9.2%) of remaining trials. Ultimately, 1576 trials (range: 1469–1611) were included for analysis.

#### Statistical analyses of experimental results

For statistical significance testing, we applied repeated-measures ANOVAs, as implemented in R package ‘ez’ (Lawrence, 2016), to test saccade metrics and timings (e.g., Figure 2), dependent variables proportion correct (Experiments 1 and 2), as well as motion-filter length and proportion filter adjusted (Experiment 3). In case of multiple factor levels and a significant Mauchly’s test for sphericity, Greenhouse-Geisser corrected p-values (*p*_*GG*_) are reported. To compute within-subject standard errors of the mean, the method of Cousineau (2005) was used. We further used linear mixed-effects models for univariate and multivariate regressions included in the ‘lme4’ R package (Bates, Mächler, Bolker, & Walker, 2015). In all models random intercepts and slopes were included for individual observers. Surfaces shown in Figure 4d and Figure 4e were fitted using mixed-effects Generalized Additive Models (GAMs), as implemented in the ‘mgcv’ package (Wood, 2017), with identity and logit link functions, respectively. The models included two orthogonal predictors − saccade direction (9 knots) and display offset relative to saccade offset (5 knots) – as thin-plate spline functions (Wood, 2003) and random smooths for each observer. A markdown document containing all analyses presented in this paper can be found at https://osf.io/ravbc/.

#### Reverse Regression

As a first step, spectral power was computed for every single stimulus image: A zero-padded, two-dimensional FFT (of three times the size of the original image in order to increase resolution) was performed and log-power was extracted using orientation-SF bandpass filters, which were applied at 18 linearly spaced orientations (bandwidth: ±5 degrees) and 20 logarithmically spaced SFs (bandwidth: ±15% of the respective SF). Mixed-effects logistic regressions (allowing random intercepts and slopes for each individual participant) were fitted separately for each display duration and orientation-SF component to predict correct trial responses in a given trial from logpower present in that same trial (Figure 3f). To make resulting beta-weights more comparable across components and display durations, log-power was standardized − weights therefore indicate log odds ratios per SD of the underlying log-power distribution. Finally, weights were fitted using an ANOVA-type GAM (Wood, 2017). It included log-SF and display duration as cubic splines (14 and 6 knots, respectively), orientation as cyclic cubic splines (13 knots), and all of their tensor product interactions. GAMs fitted on replay data had the same setup but included a treatment-coded categorical by-variable for each model term (reference: motion-only, difference: motion + suppression), in order to produce different smooths for replay conditions.

### Modeling

#### Spatiotemporal model

The model simulated the luminance modulation a single receptive field (RF) location is exposed to as it is moved across natural scenes. RFs were modeled as a quadrature pair of Gabor filters with 0^*°*^ and 90^*°*^ phase and a certain SF and orientation – an adequate model of simple receptive fields in primary visual cortex (Jones & Palmer, 1987). Nine SFs were used, spaced logarithmically from 0.05 to 5 cycles per degree (cpd), as well as nine orientations which were selected equally spaced around saccade direction. Gaussian aperture SD was determined by the SF of each filter according to a log-linear relationship between SF and RF width with a slope of -0.5 and an intercept of 0.8, as described by Anderson and Burr (1987). In order to compute Gabor filter responses to low-level image content, the stimulus image used in each trial was converted to grayscale, and the recorded eye position trajectory of the same trial was upsampled to 12 kHz, in order to be able to cover a wide range of temporal frequencies. Subsequently, for all eye position samples, Gabor filters were convolved with the current input, that is, the subset of the stimulus image defined by eye position assuming a RF eccentricity of zero. To remove baseline differences due to background luminance and varying RF size, Gabor filter responses to the mean value of the current input were subtracted. Current input was extracted from the stimulus using bilinear interpolation to achieve continuity on the sub-pixel level. Note that the responses of the pair of Gabor filters were not squared but simply summed to allow for negative values and thereby more accurately describe temporal frequencies. To extract the power in the TF domain, spectral analysis was performed on the time series (see examples in Figure 5a) using the ‘multitaper’ R package (Rahim, Burr, & Thomson, 2014). To control for the inherent SF-orientation power distribution present in the stimulus images, spectral densities in each trial, SF, and orientation were normalized by dividing values by their respective sum (Figure 5b). Prior to summing spectral power across TFs, individual TFs were weighted by human contrast sensitivity (see Contrast sensitivity filters) – that way, only power in the range of TFs that are actually resolvable by the human visual system was considered in aggregates (cf. Mostofi et al., 2020). To fit orientation tuning functions (Figure 5g), the Gaussian function

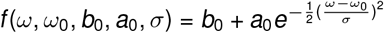

was used, where *ω* is orientation, *ω*_0_ is peak orientation, *b*_0_ and *a*_0_ are baseline and amplitude, respectively, and *σ* is tuning width. Note that only SF of ≥ 0.2 cpd entered this analysis, as only those SFs would show a significant time course. SF tuning (Figure 5e-f) was fitted with a mixed-effects GAM using thin-plate spline functions (Wood, 2003) to predict log-power based on log-SF (7 knots), display offset relative to saccade offset (4 knots), and their interaction, as well as their tensor product interaction. Random smooths were added for each observer.

#### Contrast sensitivity filters

Human contrast sensitivity, as shown in Figure 5c (cf. Kelly, 1979, Fig. 15), was modeled using the sensitivity function described in Eq. 8 in Kelly (1979):

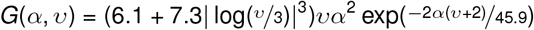

>where *α* = *·* 2*π* SF and *υ* is the current velocity of the grating that may also be defined as *υ* = ^TF^ . Note that this function was estimated in experiments using retinal stabilization of stimuli, thereby largely controlling for the influence of small eye movements on contrast sensitivity (e.g., Casile, Victor, & Rucci, 2019). The extension of the model described in Figure 6a follows the idea that if a global scene moves across the retina, then, when seen through an aperture by a RF, different orientations will move at different speeds: Orientations orthogonal to the global motion direction will move at exactly the same velocity as the global scene, whereas orientations perfectly parallel to global motion direction will not move at all. This idea is captured by the formula (cf. Scarfe & Johnston, 2011, Eq. 1)

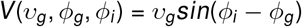

where the retinal velocity of an individual orientation is a function of the global velocity *υ*_*g*_ and the difference between that orientation *ϕ*_*i*_ and the global motion direction *ϕ*_*g*_. As saccades never move in a straight line but always exhibit some degree of curvature, we considered two velocity components, namely, velocity in the saccade’s direction *υ*_*par*_ and velocity orthogonal to the saccade’s direction *υ*_*orth*_. These velocity components can be computed after rotating saccade trajectories in a way that onset position (*x*_*on*_, *y*_*on*_) and offset position of the saccade (*x*_*off*_, *y*_*off*_) coincide with the x-axis, that is, by rotating by the saccade’s direction *α* = atan2(*y*_*off*_ − *y*_*on*_, *x*_*off*_ − *x*_*on*_). Subsequently, individual saccade samples can be adjusted relative to saccade onset position:

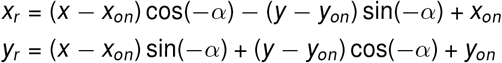

Clearly, velocities of zero could occur if *ϕ*_*i*_ = *ϕ*_*g*_, which would be incompatible with Kelly’s contrast sensitivity model. However, as saccades always introduce some retinal velocity orthogonal to their direction (Figure 6b), individual velocities of zero for a given orientation *υ*_*i*_ are extremely unlikely, if defined as:

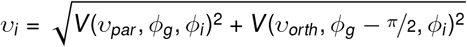

#### Saccadic suppression

The time course of saccadic suppression was based on results by Diamond et al. (2000). Data from saccadic conditions was manually digitized from Figure 2 in Ross et al. (2001a), averaged across observers MCM and MRD, and subsequently fitted with a two-piece Gaussian function to describe suppression magnitude *a*_0_ (relative to a baseline *b*) over time relative to the onset of the saccade *t*_0_:

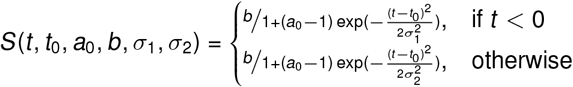

This function was well capable of capturing the asymmetric time course of saccadic suppression in the presence of both uniform (*b* = 13, *t*_0_ = − 4, *a*_0_ = 6.5, *σ*_1_ = 13, *σ*_2_ = 38) and patterned (*b* = 12, *t*_0_ = − 4, *a*_0_ = 10.9, *σ*_1_ = 20, *σ*_2_ = 50) backgrounds (Figure A4a). To model saccadic suppression in the context of natural scenes, we used the function fitted on patterned-background data for all analyses. To describe the magnitude of saccadic suppression across different SFs, we manually digitized saccade-to-fixation threshold ratios of observer D.B. from Figure 3 in Burr et al. (1982) and subsequently fitted the data with a log-linear function shown in Figure A4b:

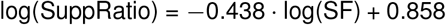

Note that as SFs increase this function predicts saccade-to-fixation ratios below 1 at approximately 7 cpd, which would indicate an enhancement of high-SF information around saccades. There is recent evidence of such enhancement at high SFs in monkeys (Niemeyer et al., 2022). Finally, these two functions were combined – under the simple assumption that suppression time courses are largely similar across SFs – to predict the magnitude of saccadic suppression across SFs and time around the saccade (Figure 5e), as suppression magnitude *a*_0_ could be predicted by SF. That is to say that fitted suppression magnitude based on the data presented by Diamond et al. (2000) was well compatible with the estimates provided by Burr et al. (1982) (see Figure A4b), considering that the former study used a background luminance of 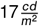, whereas the latter used 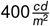 − it has been shown nu-– merous times that both increased background luminance (Burr et al., 1982; Mitrani & Yakimoff, 1971; Richards, 1969) and patterned backgrounds (Brooks & Fuchs, 1975; Chekaluk & Llewellyn, 1990; Diamond et al., 2000; Idrees et al., 2020) enhance the magnitude of suppression.

#### Motion filters

Linear motion filters (Figure 7b, left) were created using the algorithm implemented in Matlab’s ‘fspecial’ function, which was translated to the programming language R. Saccade-trajectory dependent motion filters (Figure 7b, right) were created using a custom iterative procedure based on time-resolved x-y gaze position data. As a first step, the filter kernel was allocated as a pixel matrix with dimensions specified by the range of available gaze positions. A 2D meshgrid covering the matrix’ integer x and y pixel ranges was used to locate points specified by x-y gaze positions within that matrix. By iterating over all available gaze positions, a maximum of four points within the matrix were selected that were closest to the current gaze position. These points were assigned weights that summed to the time-dependent weight (see below and Figure A5b) and were inversely proportional to their Euclidean distance to the current gaze position. At each iteration these weights were added to the filter kernel matrix. Finally, the resulting filter kernel matrix was divided by the sum of their values.

Time-dependent weights were estimated using a decay model inspired by the temporal dynamics of simple cells in primary visual cortex (Figure A5). Early visual temporal response functions were modeled using the modified gamma distribution function

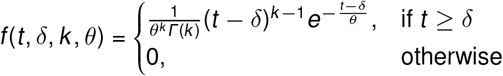

where Γ is the gamma function,δ is the response latency, *k* and *θ* are the shape and scale parameters (which must both be positive values), respectively. To set the resulting gamma shape to a certain response magnitude, all function values were divided by their respective maximum value and then multiplied by the specified amplitude *a* to return time-resolved response *R*:

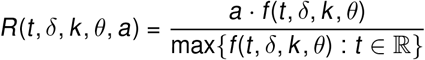

Post-stimulus spike counts (Reich et al., 2001, Fig. 2) were fitted with the above function in a mixed-effects framework that allowed *a* andδ to vary between contrast levels but not *θ* and *k* . Results revealed that the response profile could be reasonably well approximated using shape and scale parameters amounting to *k* = 1.72 and *θ* = 20.94 (Figure A5a). Finally, as illustrated in Figure A5b, we assumed that if consecutive stimuli presented at time points *t*_*i*_ elicited the same temporal responses, then the strength of the response to an individual stimulus at a target time *t*_*p*_ would be

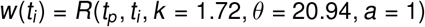

Here, we set *t*_*p*_ to the time when the response to the final stimulus reached its peak, i.e., a value of 1. This model resulted in a nonlinear but monotonic decay, that systematically downweights less recent stimulation (Figure A5c).

All modeling code can be found at https://github.com/richardschweitzer/MotionSmearModelingPlayground.

#### Estimation of perceptual consequences of postsaccadic oscillations

The amplitude of post-saccadic oscillations (PSOs) was computed as the difference between the maximum and minimum sample distance (in dva) away from the saccade onset position within the time interval of the PSO (i.e., from PSO onset to PSO offset, see Figure 2a).

To estimate the perceptual consequences that arise from these PSO amplitudes, we devised a conversion based on previous findings and theoretical considerations. First, in video-based eye tracking using the Eyelink 1000+ system, the PSO arises from the relative motion of pupil and corneal reflection. To approximate the motion of the crystalline lens, whose angular position determines the amount of shift on the retina (Deubel & Bridgeman, 1995a), we extracted relevant data from synchronous measurements of pupil and lens position (Tabernero & Artal, 2014, their Figure 2) and found that PSOs measured from the fourth Purkinje image, that is, the reflection from the back surface of the lens, were approximately 3.5 times as large as PSOs measured from the pupil center. Second, as these measurements were made with 9-degree saccades, resulting PSO amplitudes would have to be translated to those elicited by 15-degree saccades, that is, the mean amplitude in our Experiments 1 and 2 (Figure 2d). We estimated that, according to a biologically plausible model that is capable of fitting accurately families of saccades measured with video-based eye tracking (Bouzat, Freije, Frapiccini, & Gasaneo, 2018, their Figure 3b), 15-degree saccades (0.47 deg) should produce PSOs approximately 67% the amplitude of 9-degree saccades (0.7 deg). Finally, Deubel and Bridgeman (1995b) measured that one degree of PSO amplitude, when measured with a dual-Purkinje-image eye tracker and instructing saccade amplitudes of 8 deg, resulted in a perceptual shift of 0.062 dva. With these insights, we could approximate the size of the post-saccadic perceptual shift *s*_*pso*_ from a given saccade with amplitude *a*_*sac*_ and PSO amplitude *a*_*pso*_ (in deg), using the formula

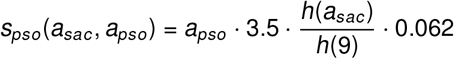

where *h*() is a nonlinear function describing the empirical relationship between *x*_*m*_ and emerging PSO amplitude shown by Bouzat et al. (2018) in their Figure 2a.

In the replay experiment, the average PSO amplitude amounted to 0.56 dva (*SD* = 0.31). Given the formula above, 15-deg saccades with the same PSO amplitude (when measured by a P-CR eye tracking system) would thus result in a perceptual error of only 0.08 dva. It can thus be approximated that the implemented replay, by displaying the measured PSO amplitude, overestimated the actual perceived amount of shift that had occurred during saccades by up to a factor of 7.

## Acknowledgements

We acknowledge the great help of Doruk Yiğ it Erigüç in data collection for Experiment 1, and R.S. thanks Karl Gegenfurtner for his valuable comments on the role of color in this series of experiments. We thank Eckart Zimmermann for his critical remarks on an earlier version of this manuscript, which we hope to have addressed in Experiment 4. R.S., M.D., M.R., and J.R. were funded by the Deutsche Forschungsgemeinschaft (DFG, German Research Foundation) under Germany’s Excellence Strategy – EXC 2002/1 “Science of Intelligence” – project number 390523135. R.S. was supported by the Studienstiftung des deutschen Volkes during the early stages of the study, as well as by the Italian Ministry for Universities and Research (MUR) and the European Union within the Next Generation EU framework (grant ‘T-GAZE’, CUP E73C22000480001) in the final stages of the study. M.R. has received funding from the European Research Council (ERC) under the European Union’s Horizon 2020 research and innovation programme (grant agreement No 865715) as well as from the Heisenberg Programme of the DFG (grants RO3579/8-1 and RO3579/12-1).

## Author contributions

R.S. and M.R. designed study. R.S. implemented experiments and photodiode tests. R.S. and M.D. collected and analyzed experimental data. R.S. performed modeling. R.S. drafted the manuscript, and all authors provided critical revisions.

## Appendix

### Appendix A

**Figure A1.**
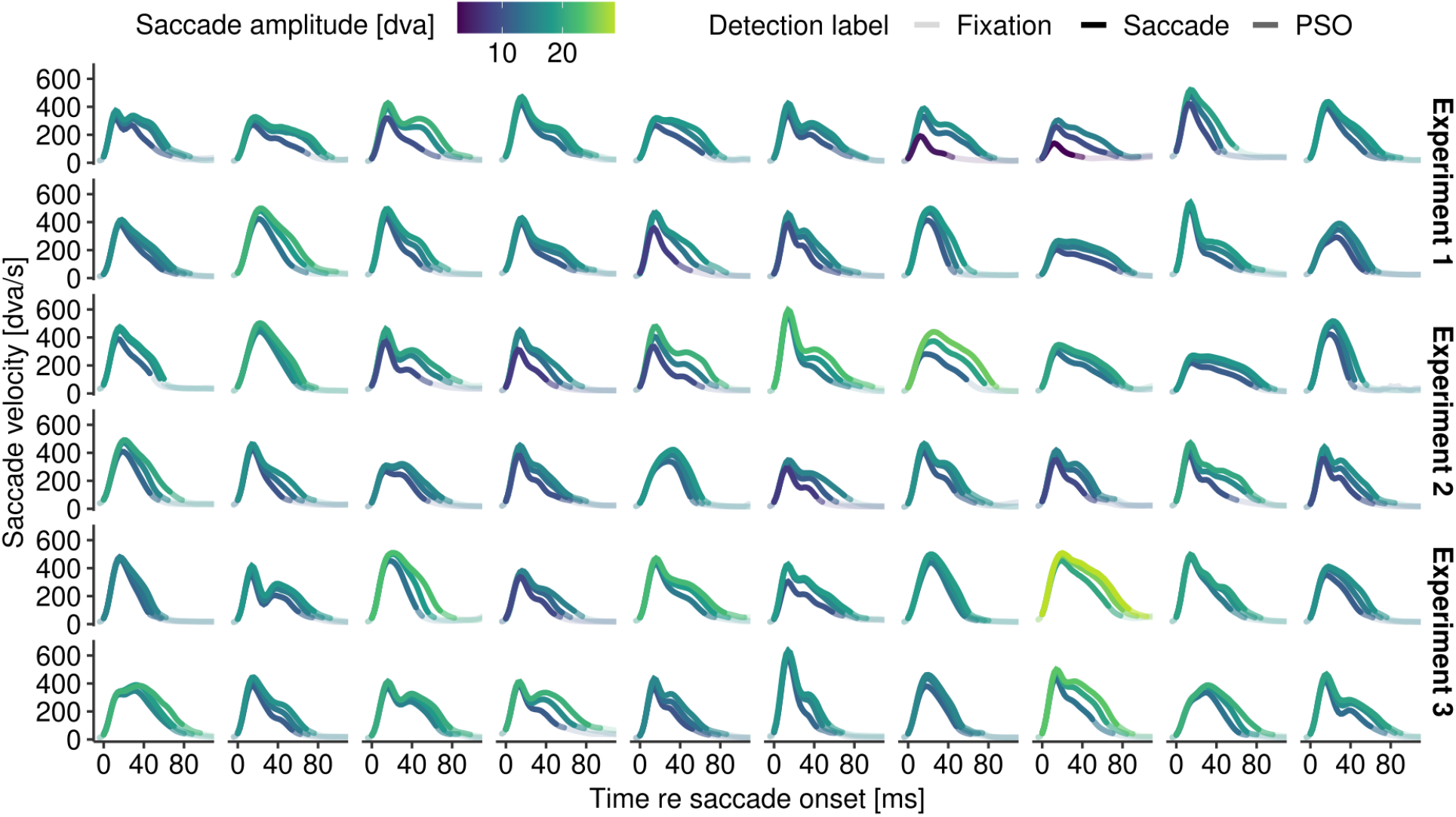
Saccadic velocity profiles of each observer. For each observer, the distribution of saccade amplitude was cut in three bins with equal an number of saccades. The average saccadic velocity profile is shown for each observer and amplitude bin. Fully opaque lines indicate samples detected as saccades, semi-transparent lines at tails of the profile indicate detected intervals of post-saccadic oscillations (PSO).

**Figure A2.**
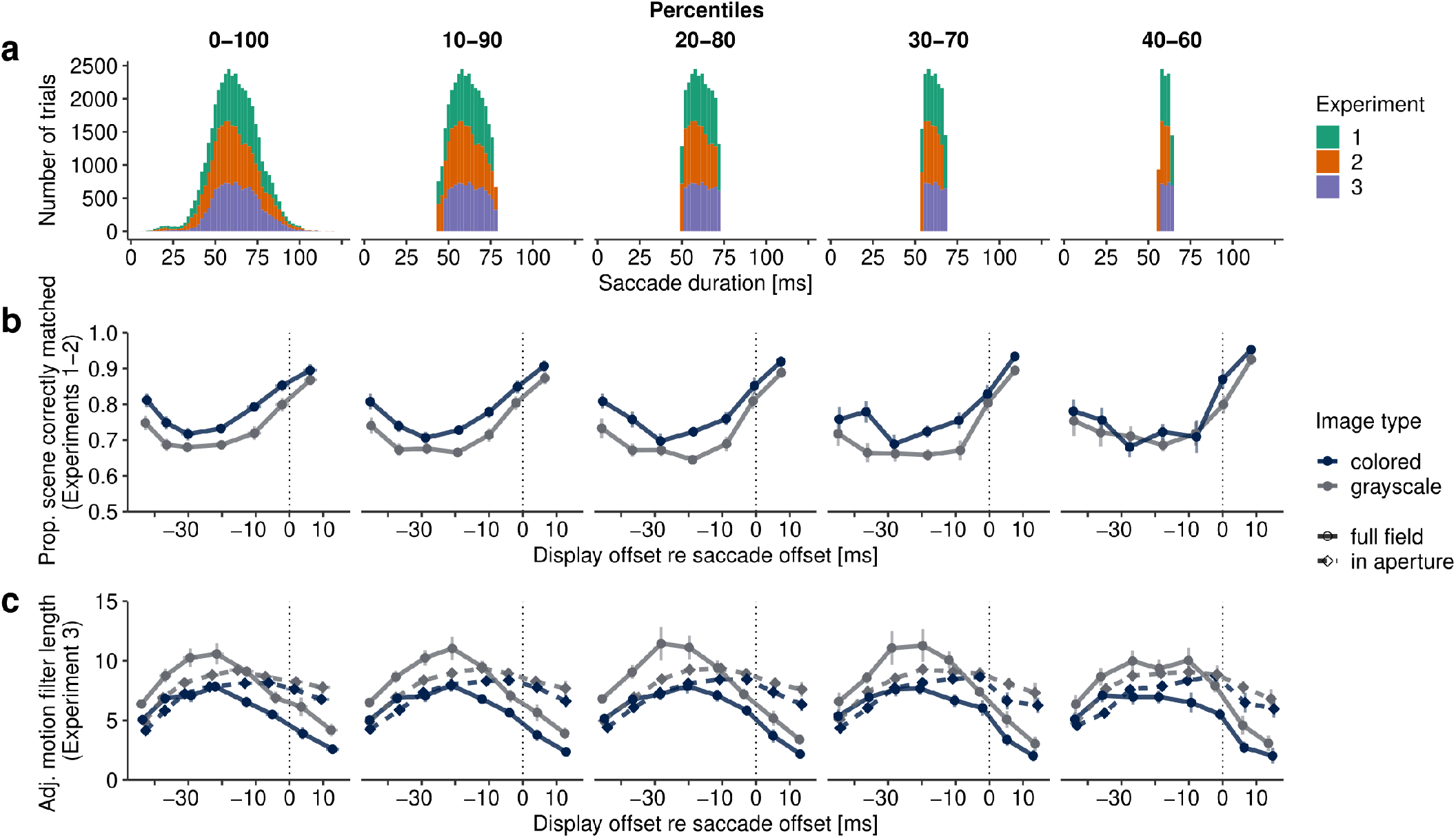
The impact of saccade-duration variance on the measured time courses. **a** Empirical cumulative distribution functions were fitted to saccade durations of each experiment. Based on percentiles of these distributions trials were systematically excluded to investigate the effect of variance in saccade duration on the estimated time courses. **b** Results of Experiments 1 and 2, averaged across experiments but otherwise as shown in Figure 3b, for different distribution widths shown in panel a. **c** Results of Experiment 3 shown in Figure 4a for different distribution widths.

**Figure A3.**
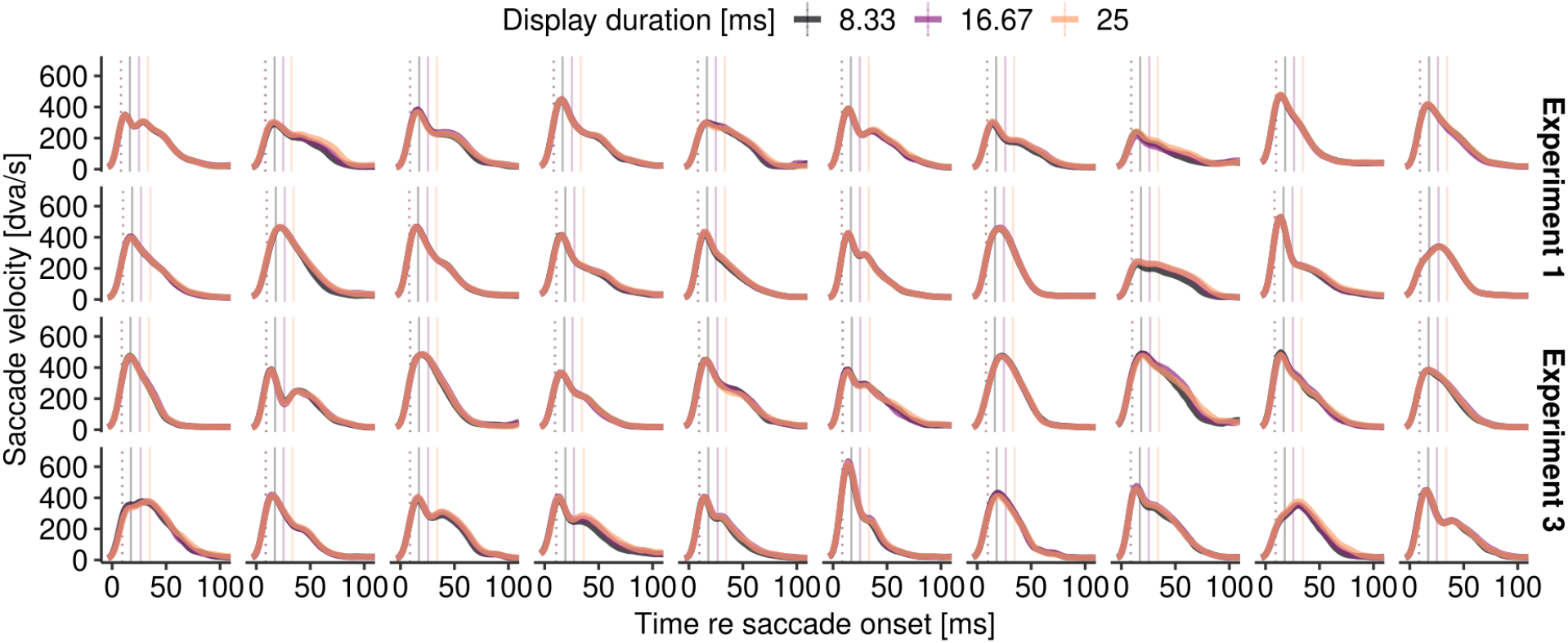
Average saccadic velocity profiles of observers in Experiments 1 and 3, plotted separately for the first three display durations 8.33, 16.67, and 25 ms. Dotted vertical lines indicate the average time of display onset, solid lines indicate average time display offset for each display duration.

**Figure A4.**
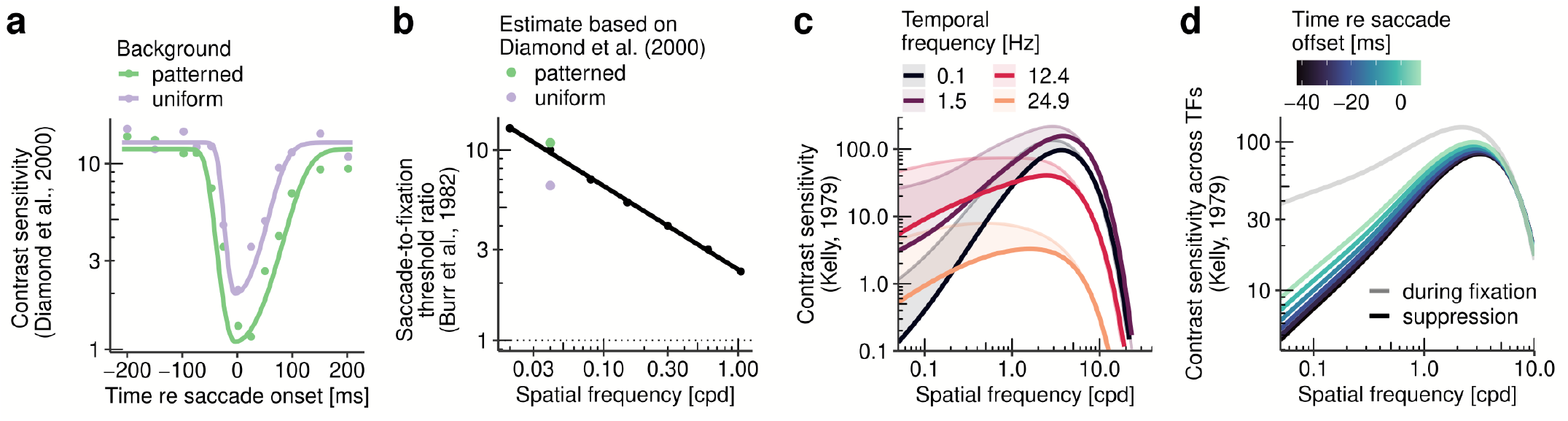
Descriptive modeling of saccadic suppression. **a** Reduction of contrast sensitivity to 0.04-cpd gratings around the onset of saccades in the presence of uniform and patterned backgrounds (data taken from Diamond et al., 2000). Data was fitted by a two-piece Gaussian function (see Saccadic suppression). **b** Dependence of saccadic suppression magnitude (i.e., ratio of detection thresholds during saccades and fixation) on spatial frequency of the displayed stimulus (data taken from Burr et al., 1982), approximated by a log-linear function (see Saccadic suppression). Estimates from panel a are shown as colored dots. **c** Estimated effect of saccadic suppression around 7 ms after saccade onset on the human contrast sensitivity profile (Kelly, 1979). Shaded areas indicate the reduction of contrast sensitivity (thin lines: baseline, thick lines: after saccadic suppression) for four different temporal frequencies. **d** Effect of saccadic suppression as a function of time relative to saccade offset. All contrast sensitivity curves (gray line: baseline) show average sensitivity across temporal frequencies (1–50 Hz).

**Figure A5.**
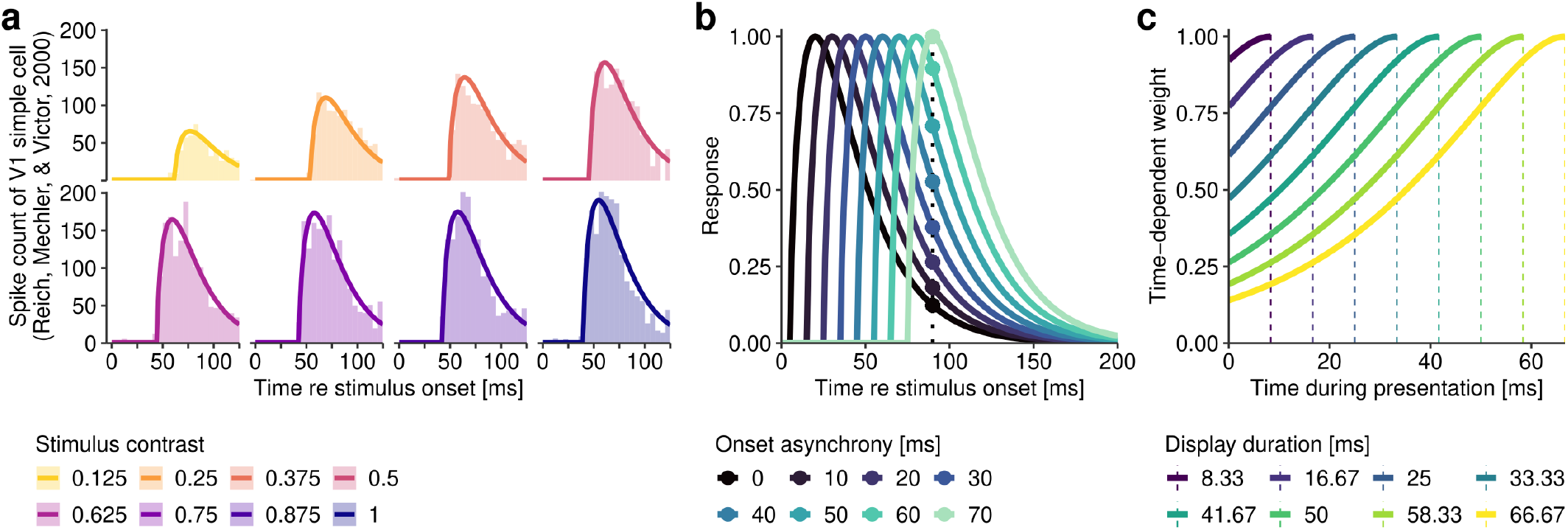
Model of visual decay. **a** Temporal response functions of a V1 simple cell. Post-stimulus time spike-count histograms (recorded for stationary gratings of eight different contrast levels) were manually digitized from figure* 2 in Reich et al. (2001), arranged in 5-ms time bins, and fitted by an extended gamma distribution function with fixed shape and scale parameters across all stimulus contrasts (see Motion filters). **b** Normalized temporal responses (using the function fitted in panel a, but for illustration purposes assuming an unrealistic latency of 5 ms) to subsequent stimulation at varying onset asynchronies (0–70 ms). Vertical dashed line indicates the time of the maximum most recent response. The response magnitude at that time point determine respective time-dependent weights (highlighted dots). **c** Time-dependent weights computed for all display durations used in experiments. Dashed lines indicate display offset and therefore most recent sample (by default with a value of 1).

**Figure A6.**
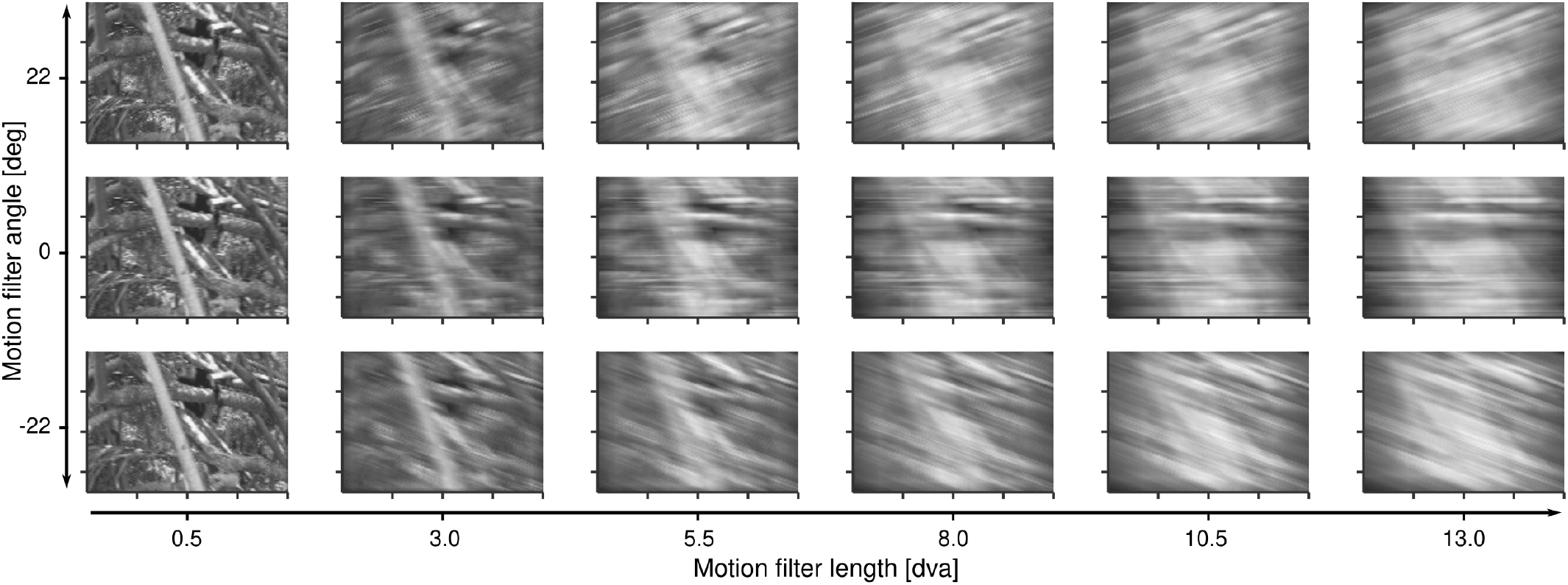
Examples of motion filter output. Illustration of the effect of applying linear motion filters with different angles (panel rows) and lengths (panel columns).

**Figure A7.**
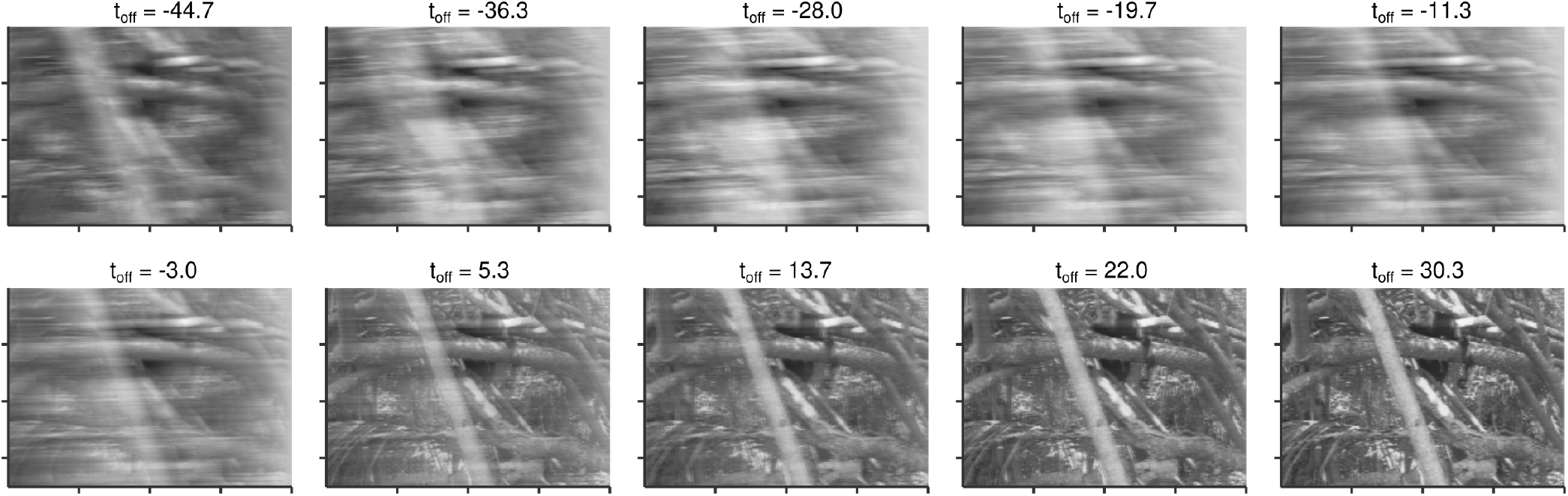
Examples of trajectory-dependent motion filter output. Predicted motion blur for different display offsets around saccade offset (t_off_). All filters were estimated based on the average saccade shown in Figure 6b. The last two panels (right, bottom row) show output for two additional display durations, that is, 75.0 and 83.3 ms, respectively, which were not used in experiments. These model predictions assumed a temporal response described by k = 1.72 and θ = 20.94 (for details, see Motion filters).

## Notes

We have no conflict of interest to disclose.

https://osf.io/bf246/

https://osf.io/ue6cd/

https://osf.io/ravbc/

https://osf.io/56mqg/

https://github.com/richardschweitzer/LM03_lightmeter

https://github.com/richardschweitzer/MotionSmearModelingPlayground

### Competing Interest Statement

The authors have declared no competing interest.

### Summary of Updates

Added Experiment 4 in Methods and Results, as well as a significant extension of the Discussion.

## References

Adams, W. J., Elder, J. H., Graf, E. W., Leyland, J., Lugtigheid, A. J., & Muryy, A. (2016). The Southampton-York natural scenes (SYNS) dataset: Statistics of surface attitude. Scientific reports, 6, 35805.

Adelson, E. H., & Bergen, J. R. (1985, February). Spatiotemporal energy models for the perception of motion. J. Opt. Soc. Am. A, 2(2), 284–299.

Alais, D., Apthorp, D., Karmann, A., & Cass, J. (2011). Temporal integration of movement: the time-course of motion streaks revealed by masking. PloS one, 6(12).

Albrecht, D. G., Geisler, W. S., Frazor, R. A., & Crane, A. M. (2002). Visual cortex neurons of monkeys and cats: temporal dynamics of the contrast response function. Journal of neurophysiology, 88(2), 888–913.

Anderson, S. J., & Burr, D. C. (1987). Receptive field size of human motion detection units. Vision research, 27 (4), 621–635.

Apthorp, D., Cass, J., & Alais, D. (2010). Orientation tuning of contrast masking caused by motion streaks. Journal of Vision, 10(10), 11.

Apthorp, D., Cass, J., & Alais, D. (2011). The spatial tuning of “motion streak” mechanisms revealed by masking and adaptation. Journal of Vision, 11(7), 17.

Apthorp, D., Schwarzkopf, D. S., Kaul, C., Bahrami, B., Alais, D., & Rees, G. (2013). Direct evidence for encoding of motion streaks in human visual cortex. Proceedings of the Royal Society B: Biological Sciences, 280(1752), 20122339.

Bahill, A. T., Clark, M. R., & Stark, L. (1975, January). The main sequence, a tool for studying human eye movements. Mathematical Biosciences, 24(3), 191–204.

Baker, D. H., Vilidaite, G., Lygo, F. A., Smith, A. K., Flack, T. R., Gouws, A. D., & Andrews, T. J. (2021). Power contours: Optimising sample size and precision in experimental psychology and human neuroscience. Psychological methods, 26(3), 295.

Balsdon, T., Schweitzer, R., Watson, T. L., & Rolfs, M. (2018). All is not lost: Post-saccadic contributions to the perceptual omission of intra-saccadic streaks. Consciousness and cognition, 64, 19–31.

Bates, D., Mächler, M., Bolker, B., & Walker, S. (2015). Fitting linear mixed-effects models using lme4. Journal of Statistical Software, 67 (1), 1–48.

Becker, W. (1989). The neurobiology of saccadic eye movements. metrics. Reviews of oculomotor research, 3, 13.

Bedell, H. E., & Yang, J. (2001). The attenuation of perceived image smear during saccades. Vision Research, 41(4), 521–528.

Binda, P., & Morrone, M. C. (2018). Vision during saccadic eye movements. Annual review of vision science, 4, 193–213.

Binda, P., & Morrone, M. C. (2022). Vision: Optimizing each glimpse. Current Biology, 32(12), R567–R569.

Boi, M., Poletti, M., Victor, J. D., & Rucci, M. (2017). Consequences of the oculomotor cycle for the dynamics of perception. Current Biology, 27 (9), 1268–1277.

Bouzat, S., Freije, M. L., Frapiccini, A. L., & Gasaneo, G. (2018). Inertial movements of the iris as the origin of postsaccadic oscillations. Physical review letters, 120(17), 178101.

Braun, D. I., Schütz, A. C., & Gegenfurtner, K. R. (2017, July). Visual sensitivity for luminance and chromatic stimuli during the execution of smooth pursuit and saccadic eye movements. Vision Research, 136, 57–69.

Breitmeyer, B. G., & Ganz, L. (1976). Implications of sustained and transient channels for theories of visual pattern masking, saccadic suppression, and information processing. Psychological review, 83(1), 1–36.

Bremmer, F., Kubischik, M., Hoffmann, K.-P., & Krekelberg, B. (2009). Neural dynamics of saccadic suppression. Journal of Neuroscience, 29(40), 12374–12383. doi: 10.1523/JNEUROSCI.2908-09.2009

Bridgeman, B., Van der Heijden, A. H. C., & Velichkovsky, B. M. (1994). A theory of visual stability across saccadic eye movements. Behavioral and Brain Sciences, 17 (2), 247–258.

Brooks, B. A., & Fuchs, A. F. (1975). Influence of stimulus parameters on visual sensitivity during saccadic eye movement. Vision Research, 15(12), 1389–1398.

Brooks, B. A., Impelman, D., & Lum, J. (1981). Backward and forward masking associated with saccadic eye movement. Perception & Psychophysics, 30(1), 62–70.

Burr, D. (1980). Motion smear. Nature, 284(5752), 164–165.

Burr, D. (1981). Temporal summation of moving images by the human visual system. Proc. R. Soc. Lond. B, 211(1184), 321–339.

Burr, D., Holt, J., Johnstone, J., & Ross, J. (1982). Selective depression of motion sensitivity during saccades. The Journal of physiology, 333(1), 1–15.

Burr, D., Morrone, M., & Ross, J. (1994). Selective suppression of the magnocellular visual pathway during saccadic eye movements. Nature, 371(6497), 511–513.

Burr, D., & Ross, J. (1982). Contrast sensitivity at high velocities. Vision research, 22(4), 479–484.

Campbell, F., & Wurtz, R. (1978). Saccadic omission: why we do not see a grey-out during a saccadic eye movement. Vision research, 18(10), 1297–1303.

Casile, A., Victor, J. D., & Rucci, M. (2019). Contrast sensitivity reveals an oculomotor strategy for temporally encoding space. Elife, 8, e40924.

Castet, E. (2010). Perception of intra-saccadic motion. In Dynamics of visual motion processing, chapter 10 (pp. 213–238). Springer.

Castet, E., Jeanjean, S., & Masson, G. S. (2001). “Saccadic suppression” - no need for an active extra-retinal mechanism. Trends in Neurosciences, 24(6), 316–317.

Castet, E., Jeanjean, S., & Masson, G. S. (2002). Motion perception of saccade-induced retinal translation. Proceedings of the National Academy of Sciences, 99(23), 15159–15163.

Castet, E., & Masson, G. (2000). Motion perception during saccadic eye movements. Nature Neuroscience, 2, 177–183.

Cavanagh, P., Hunt, A. R., Afraz, A., & Rolfs, M. (2010). Visual stability based on remapping of attention pointers. Trends in cognitive sciences, 14(4), 147–153.

Chekaluk, E., & Llewellyn, K. R. (1990). Visual stimulus input, saccadic suppression, and detection of information from the postsaccade scene. Perception & Psychophysics, 48(2), 135–142.

Coppola, D. M., Purves, H. R., McCoy, A. N., & Purves, D. (1998). The distribution of oriented contours in the real world. Proceedings of the National Academy of Sciences, 95(7), 4002–4006.

Cornelisen, F., Peters, E., & Palmer, J. (2002). The eyelink toolbox: eyetracking with matlab and the psychophysics toolbox. Behaviour Research Methods, 34(4), 613–617.

Cousineau, D. (2005). Confidence intervals in within-subject designs: A simpler solution to Loftus and Masson’s method. Tutorials in quantitative methods for psychology, 1(1), 42–45.

Dassonville, P., Schlag, J., & Schlag-Rey, M. (1992). Oculomotor localization relies on a damped representation of saccadic eye displacement in human and nonhuman primates. Visual neuroscience, 9(3-4), 261–269.

Denagamage, S., Morton, M. P., Hudson, N. V., Reynolds, J. H., Jadi, M. P., & Nandy, A. S. (2023). Laminar mechanisms of saccadic suppression in primate visual cortex. Cell reports, 42(7).

Deubel, H., & Bridgeman, B. (1995a). Fourth purkinje image signals reveal eye-lens deviations and retinal image distortions during saccades. Vision research, 35(4), 529–538.

Deubel, H., & Bridgeman, B. (1995b). Perceptual consequences of ocular lens overshoot during saccadic eye movements. Vision Research, 35(20), 2897–2902.

Diamond, M., Ross, J., & Morrone, M. (2000). Extraretinal control of saccadic suppression. Journal of Neuroscience, 20(9), 3449–3455.

Dong, D. W., & Atick, J. J. (1995). Statistics of natural time-varying images. Network: Computation in Neural Systems, 6(3), 345.

Duyck, M., Collins, T., & Wexler, M. (2016). Masking the saccadic smear. Journal of vision, 16(10), 1.

Duyck, M., Collins, T., & Wexler, M. (2021). Visual continuity during blinks and alterations in time perception. Journal of Experimental Psychology: Human Perception and Performance, 47 (1), 1.

Duyck, M., Wexler, M., Castet, E., & Collins, T. (2018). Motion masking by stationary objects: a study of simulated saccades. i-Perception, 9(3), 2041669518773111.

Engbert, R., & Kliegl, R. (2003). Microsaccades uncover the orientation of covert attention. Vision research, 43(9), 1035–1045.

Engbert, R., & Mergenthaler, K. (2006). Microsaccades are triggered by low retinal image slip. Proceedings of the National Academy of Sciences, 103(18), 7192–7197.

Frazor, R. A., Albrecht, D. G., Geisler, W. S., & Crane, A. M. (2004). Visual cortex neurons of monkeys and cats: temporal dynamics of the spatial frequency response function. Journal of Neurophysiology, 91(6), 2607–2627.

Frost, A., & Niemeier, M. (2015). Suppression and reversal of motion perception around the time of the saccade. Frontiers in Systems Neuroscience, 9(143). doi: 10.3389/fnsys.2015.00143

García-Pérez, M. A., & Peli, E. (2001). Intrasaccadic perception. Journal of Neuroscience, 21(18), 7313–7322.

García-Pérez, M. A., & Peli, E. (2011). Visual contrast processing is largely unaltered during saccades. Frontiers in Psychology, 2, –. doi: 10.3389/fpsyg.2011.00247

Gegenfurtner, K. R., & Rieger, J. (2000). Sensory and cognitive contributions of color to the recognition of natural scenes. Current Biology, 10(13), 805–808.

Geisler, W. S. (1999). Motion streaks provide a spatial code for motion direction. Nature, 400(6739), 65–69.

Geisler, W. S., Albrecht, D. G., Crane, A. M., & Stern, L. (2001). Motion direction signals in the primary visual cortex of cat and monkey. Visual neuroscience, 18(4), 501–516.

Hansen, T., & Gegenfurtner, K. R. (2009). Independence of color and luminance edges in natural scenes. Visual neuroscience, 26(1), 35–49.

Hershberger, W. A., & Jordan, J. S. (1998, January). The phantom array: A perisaccadic illusion of visual direction. The Psychological Record, 48(1), 21–32.

Hooge, I., Holmqvist, K., & Nyström, M. (2016). The pupil is faster than the corneal reflection (cr): Are video based pupil-cr eye trackers suitable for studying detailed dynamics of eye movements? Vision research, 128, 6–18.

Ibbotson, M. R., & Cloherty, S. L. (2009, June). Visual perception: Saccadic omission - suppression or temporal masking? Current Biology, 19(12), 493–496.

Ibbotson, M. R., Crowder, N. A., Cloherty, S. L., Price, N. S. C., & Mustari, M. J. (2008). Saccadic modulation of neural responses: Possible roles in saccadic suppression, enhancement, and time compression. The Journal of Neuroscience, 28(43), 10952–10960. doi: 10.1523/JNEUROSCI.3950-08.2008

Ibbotson, M. R., Price, N. S., Crowder, N. A., Ono, S., & Mustari, M. J. (2007). Enhanced motion sensitivity follows saccadic suppression in the superior temporal sulcus of the macaque cortex. Cerebral cortex, 17 (5), 1129–1138.

Idrees, S., Baumann, M. P., Franke, F., Münch, T. A., & Hafed, Z. M. (2020). Perceptual saccadic suppression starts in the retina. Nature Communications, 11(1), 1–19.

Ilg, U. J., Bridgeman, B., & Hoffmann, K. P. (1989). Influence of mechanical disturbance on oculomotor behavior. Vision research, 29(5), 545–551.

Jancke, D. (2000). Orientation formed by a spot’s trajectory: a two-dimensional population approach in primary visual cortex. Journal of Neuroscience, 20(14), RC86–RC86.

Jones, J. P., & Palmer, L. A. (1987). An evaluation of the two-dimensional gabor filter model of simple receptive fields in cat striate cortex. Journal of neurophysiology, 58(6), 1233–1258.

Jordan, J. S., & Hershberger, W. A. (1994). Timing the shift in retinal local signs that accompanies a saccadic eye movement. Perception & Psychophysics, 55, 657–666.

Kelly, D. H. (1975). Luminous and chromatic flickering patterns have opposite effects. Science, 188(4186), 371–372.

Kelly, D. H. (1977). Visual contrast sensitivity. Optica Acta: International Journal of Optics, 24(2), 107–129.

Kelly, D. H. (1979). Motion and vision. ii. stabilized spatio-temporal threshold surface. J. Opt. Soc. Am., 69(10), 1340–1349.

Kleiner, M., Brainard, D., Pelli, D., Ingling, A., Murray, R., & Broussard, C. (2007). What is new in psychtoolbox-3. Perception, 36(14), 1–16.

Knöll, J., Binda, P., Morrone, M. C., & Bremmer, F. (2011). Spatiotemporal profile of peri-saccadic contrast sensitivity. Journal of vision, 11(14), 15.

Komban, S. J., Kremkow, J., Jin, J., Wang, Y., Lashgari, R., Li, X., … Alonso, J.-M. (2014). Neuronal and perceptual differences in the temporal processing of darks and lights. Neuron, 82(1), 224–234.

Latour, P. L. (1962). Visual threshold during eye movements. Vision Research, 2(3), 261–262.

Lawrence, M. A. (2016). ez: Easy analysis and visualization of factorial experiments [Computer software manual]. Retrieved from https://CRAN.R-project.org/package=ez (R package version 4. 4-0)

Lebedev, S., Van Gelder, P., & Tsui, W. H. (1996). Square-root relations between main saccadic parameters. Investigative Ophthalmology & Visual Science, 37 (13), 2750–2758.

MacKay, D. M. (1973). Visual stability and voluntary eye movements. In Central processing of visual information a: integrative functions and comparative data (pp. 307–331). Springer.

Martinez-Conde, S., Macknik, S. L., & Hubel, D. H. (2004, March). The role of fixational eye movements in visual perception. Nature Reviews Neuroscience, 5(3), 229–240.

Masselink, J., & Lappe, M. (2021). Visuomotor learning from postdictive motor error. Elife, 10, e64278.

Mateeff, S. (1978). Saccadic eye movements and localization of visual stimuli. Perception & Psychophysics, 24(3), 215–224.

Matin, E. (1974). Saccadic suppression: a review and an analysis. Psychological bulletin, 81(12), 899.

Matin, E., Clymer, A., & Matin, L. (1972). Metacontrast and saccadic suppression. Science, 178(4057), 179–182.

Matin, L., & Pearce, D. G. (1965). Visual perception of direction for stimuli flashed during voluntary saccadic eye movements. Science, 148(3676), 1485–1488.

Mitrani, L., & Yakimoff, N. (1970). Smearing of the retinal image during voluntary saccadic eye movements. Vision Research, 10(5), 405–409.

Mitrani, L., & Yakimoff, N. (1971). Is saccadic suppression really saccadic? Vision Research, 11(10), 1157–1161.

Mitrani, L., Yakimoff, N., & Mateeff, S. T. (1973). Saccadic suppression in the presence of structured background. Vision research.

Morris, A. P., Kubischik, M., Hoffmann, K.-P., Krekelberg, B., & Bremmer, F. (2012). Dynamics of eye-position signals in the dorsal visual system. Current Biology, 22(3), 173–179.

Mostofi, N., Zhao, Z., Intoy, J., Boi, M., Victor, J. D., & Rucci, M. (2020). Spatiotemporal content of saccade transients. Current Biology, 30(20), 3999–4008.

Neri, P. (2014). Semantic control of feature extraction from natural scenes. Journal of Neuroscience, 34(6), 2374–2388.

Nicolas, G., Castet, E., Rabier, A., Kristensen, E., Dojat, M., & Guérin-Dugué, A. (2021). Neural correlates of intrasaccadic motion perception. Journal of Vision, 21(11), 19–19.

Niemeyer, J. E., Akers-Campbell, S., Gregoire, A., & Paradiso, M. A. (2022). Perceptual enhancement and suppression correlate with v1 neural activity during active sensing. Current Biology .

Nishida, S., Watanabe, J., Kuriki, I., & Tokimoto, T. (2007). Human visual system integrates color signals along a motion trajectory. Current Biology, 17 (4), 366–372.

Nyström, M., & Holmqvist, K. (2010). An adaptive algorithm for fixation, saccade, and glissade detection in eyetracking data. Behavior research methods, 42(1), 188–204.

Pola, J. (2004). Models of the mechanism underlying perceived location of a perisaccadic flash. Vision research, 44(24), 2799–2813.

Pomè, A., Schlichting, N., Fritz, C., & Zimmermann, E. (2024). Prediction of sensorimotor contingencies generates saccadic omission. Current Biology .

Rahim, K. J., Burr, W. S., & Thomson, D. J. (2014). Applications of multitaper spectral analysis to nonstation-ary data (Doctoral dissertation, Queen’s University). Retrieved from https://CRAN.R-project.org/package=multitaper (R package version 1. 0–15)

Reich, D. S., Mechler, F., & Victor, J. D. (2001). Temporal coding of contrast in primary visual cortex: when, what, and why. Journal of neurophysiology, 85(3), 1039–1050.

Reichardt, W. (1987). Evaluation of optical motion information by movement detectors. Journal of Comparative Physiology A, 161(4), 533–547.

Richards, W. (1968). Visual suppression during passive eye movement. J. Opt. Soc. Am., 58(8), 1159–1160.

Richards, W. (1969). Saccadic suppression. Journal of the Optical Society of America, 59(5), 617–623.

Riggs, L. A., & Manning, K. A. (1982, July). Saccadic suppression under conditions of whiteout. Invest. Ophthalmol. Vis. Sci., 23(1), 138–143.

Robson, J. G. (1966). Spatial and temporal contrast-sensitivity functions of the visual system. Josa, 56(8), 1141–1142.

Rolfs, M. (2015). Attention in active vision: A perspective on perceptual continuity across saccades. Perception, 44, 900–919.

Rolfs, M., & Schweitzer, R. (2022). Coupling perception to action through incidental sensory consequences of motor behaviour. Nature Reviews Psychology, 1(2), 112–123.

Rolfs, M., Schweitzer, R., Castet, E., Watson, T. L., & Ohl, S. (2023). Lawful kinematics link eye movements to the limits of high-speed perception. bioRxiv, 2023–07.

Ross, J., Morrone, M., Goldberg, M., & Burr, D. (2001a). Changes in visual perception at the time of saccades. Trends in neurosciences, 24(2), 113–121.

Ross, J., Morrone, M., Goldberg, M. E., & Burr, D. C. (2001b). Response: “Saccadic suppression” - no need for an active extra-retinal mechanism. Trends in Neurosciences, 24(6), 317–318. doi: 10.1016/S0166-2236(00)01827-0

Rucci, M., Ahissar, E., & Burr, D. (2018). Temporal coding of visual space. Trends in cognitive sciences, 22(10), 883–895.

Scarfe, P., & Johnston, A. (2011). Global motion coherence can influence the representation of ambiguous local motion. Journal of Vision, 11(12), 6–6.

Schlag, J., & Schlag-Rey, M. (1995). Illusory localization of stimuli flashed in the dark before saccades. Vision research, 35(16), 2347–2357.

Scholes, C., McGraw, P. V., & Roach, N. W. (2021). Learning to silence saccadic suppression. Proceedings of the National Academy of Sciences, 118(6), e2012937118.

Schweitzer, R., & Rolfs, M. (2020a). An adaptive algorithm for fast and reliable online saccade detection. Behavior research methods, 52(3), 1122–1139. doi: 10.3758/s13428-019-01304-3

Schweitzer, R., & Rolfs, M. (2020b). Intra-saccadic motion streaks as cues to linking object locations across saccades. Journal of Vision, 20(4), 17. doi: 10.1167/jov.20.4.17

Schweitzer, R., & Rolfs, M. (2021). Intrasaccadic motion streaks jump-start gaze correction. Science Advances, 7 (30), eabf2218. doi: 10.1126/sciadv.abf2218

Schweitzer, R., & Rolfs, M. (2022). Definition, modeling, and detection of saccades in the face of post-saccadic oscillations. In S. Stuart (Ed.), Eye tracking: Background, methods, and applications (pp. 69–95). New York, NY: Springer US. doi: 10.1007/978-1-0716-2391-6_5

Schweitzer, R., Watson, T., Watson, J., & Rolfs, M. (2019). The joy of retinal painting: A build-it-yourself device for intrasaccadic presentations. Perception, 48(10), 1020–1025. doi: 10.1177/0301006619867868

Shi, L. (2017). An evaluation of an lcd display with 240 hz frame rate for visual psychophysics experiments. i-Perception, 8(5), 2041669517736788.

Smit, A., & Van Gisbergen, J. (1990). An analysis of curvature in fast and slow human saccades. Experimental Brain Research, 81(2), 335–345.

Sperling, G., & Speelman, R. G. (1965). Visual spatial localization during object motion, apparent object motion, and image motion produced by eye movements [abstract]. Journal of the Optical Society of America, 55, 1576–1577.

Stevenson, S., Volkmann, F., Kelly, J., & Riggs, L. A. (1986). Dependence of visual suppression on the amplitudes of saccades and blinks. Vision research, 26(11), 1815–1824.

Sylvester, R., Haynes, J., & Rees, G. (2005). Saccades differentially modulate human lgn and v1 responses in the presence and absence of visual stimulation. Current Biology, 15(1), 37–41.

Tabernero, J., & Artal, P. (2014). Lens oscillations in the human eye. implications for post-saccadic suppression of vision. PloS one, 9(4), e95764.

Teichert, T., Klingenhoefer, S., Wachtler, T., & Bremmer, F. (2010). Perisaccadic mislocalization as optimal percept. Journal of vision, 10(8), 19.

Thiele, A., Henning, P., Kubischik, M., & Hoffmann, K.-P. (2002). Neural mechanisms of saccadic suppression. Science, 295(5564), 2460–2462.

Todorov, V., & Filzmoser, P. (2009). An object-oriented framework for robust multivariate analysis. Journal of Statistical Software, 32(3), 1–47.

Tolhurst, D. J., Tadmor, Y., & Chao, T. (1992). Amplitude spectra of natural images. Ophthalmic and Physiological Optics, 12(2), 229–232.

Torralba, A., & Oliva, A. (2003). Statistics of natural image categories. Network: computation in neural systems, 14(3), 391.

Van der Schaaf, v. A., & van Hateren, J. v. (1996). Modelling the power spectra of natural images: statistics and information. Vision research, 36(17), 2759–2770.

Van der Stigchel, S., Meeter, M., & Theeuwes, J. (2006). Eye movement trajectories and what they tell us. Neuroscience & biobehavioral reviews, 30(5), 666–679.

Van Opstal, A., & Van Gisbergen, J. (1987). Skewness of saccadic velocity profiles: a unifying parameter for normal and slow saccades. Vision research, 27 (5), 731–745.

Viviani, P., Berthoz, A., & Tracey, D. (1977). The curvature of oblique saccades. Vision research, 17 (5), 661–664.

Volkmann, F. C. (1986). Human visual suppression. Vision research, 26(9), 1401–1416.

Volkmann, F. C., Riggs, L., White, K., & Moore, R. (1978). Contrast sensitivity during saccadic eye movements. Vision research, 18(9), 1193–1199.

Wexler, M., & Cavanagh, P. (2019). Fast motion drags shape. Journal of Vision, 19(10), 288c–288c. (Poster at VSS)

Wexler, M., & Collins, T. (2014). Orthogonal steps relieve saccadic suppression. Journal of Vision, 14(2), 13.

Wichmann, F. A., Sharpe, L. T., & Gegenfurtner, K. R. (2002). The contributions of color to recognition memory for natural scenes. Journal of Experimental Psychology: Learning, Memory, and Cognition, 28(3), 509.

Wood, S. N. (2003). Thin plate regression splines. Journal of the Royal Statistical Society: Series B (Statistical Methodology), 65(1), 95–114.

Wood, S. N. (2017). Generalized additive models: an introduction with r. Chapman and Hall/CRC.

Wurtz, R. H. (2008). Neuronal mechanisms of visual stability. Vision research, 48(20), 2070–2089.

Wurtz, R. H. (2018). Corollary discharge contributions to perceptual continuity across saccades. Annu. Rev. Vis. Sci., 4(1), 215–237. doi: 10.1146/annurev-vision-102016-061207

Zhang, R., Isola, P., Efros, A. A., Shechtman, E., & Wang, O. (2018). The unreasonable effectiveness of deep features as a perceptual metric. In Proceedings of the ieee conference on computer vision and pattern recognition (pp. 586–595).

Zimmermann, E. (2020). Saccade suppression depends on context. Elife, 9, e49700.

Zimmermann, E., & Lange, J. (2022). Saccade suppression of displacements, but not of contrast, depends on context. Journal of Vision, 22(10), 10–10.

